# Molecular definition of distinct active zone protein machineries for Ca^2+^ channel clustering and synaptic vesicle priming

**DOI:** 10.1101/2023.10.27.564439

**Authors:** Javier Emperador-Melero, Jonathan W. Andersen, Sarah R. Metzbower, Aaron D. Levy, Poorna A. Dharmasri, Giovanni de Nola, Thomas A. Blanpied, Pascal S. Kaeser

## Abstract

Action potentials trigger neurotransmitter release with minimal delay. Active zones mediate this temporal precision by co-organizing primed vesicles with Ca_V_2 Ca^2+^ channels. The presumed model is that scaffolding proteins directly tether primed vesicles to Ca_V_2s. We find that Ca_V_2 clustering and vesicle priming are executed by separate machineries. At hippocampal synapses, Ca_V_2 nanoclusters are positioned at variable distances from those of the priming protein Munc13. The active zone organizer RIM anchors both proteins, but distinct interaction motifs independently execute these functions. In heterologous cells, Liprin-α and RIM from co- assemblies that are separate from Ca_V_2-organizing complexes upon co-transfection. At synapses, Liprin-α1-4 knockout impairs vesicle priming, but not Ca_V_2 clustering. The cell adhesion protein PTPσ recruits Liprin-α, RIM and Munc13 into priming complexes without co- clustering of Ca_V_2s. We conclude that active zones consist of distinct complexes to organize Ca_V_2s and vesicle priming, and Liprin-α and PTPσ specifically support priming site assembly.

## Introduction

The active zone is a protein scaffold that enables fast neurotransmitter release. It mediates the priming of synaptic vesicles to add them to the readily releasable pool and the clustering of voltage-gated Ca^2+^ channels of the Ca_V_2 family^1,2^. Work over the past decades has generated essential insights into how active zone functions are executed. Synaptic vesicle priming depends on the recruitment of Munc13 by the active zone organizer RIM, and Munc13 supports subsequent assembly of the exocytic SNARE complex^3–7^. The clustering of Ca_V_2s is also orchestrated by RIM, in conjunction with tripartite interactions with RIM-BP^8–12^. At least three additional protein families further sustain active zone functions. First, ELKS is important for Ca^2+^ influx and for the generation of primed vesicles^13–16^. Second, Piccolo/Bassoon are long-range tethers and Bassoon contributes to Ca_V_2 clustering via RIM-BP^17–19^. Third, Liprin-α recruits presynaptic material and may act upstream in active zone assembly^20–25^. This groundwork has led to mechanistic models for vesicle priming and Ca_V_2 clustering, but it has not been possible to define how the active zone is organized to simultaneously maintain these two functions.

The simplest model suggests that synaptic vesicle priming and Ca_V_2 clustering are executed by the same protein complex containing all active zone proteins. This is rooted in the observation that all key active zone proteins interact with one another and that most of them contribute to both functions^4,8,10–12,15,16,19,26–29^. Furthermore, Ca_V_2s themselves might directly tether vesicles^30–32^. Additional strong support for a “single complex” model comes from recent findings that vesicles can be recruited to the surface of reconstituted condensates of Ca^2+^ channel-organizing complexes consisting of RIM, RIM-BP and Ca_V_2^33^ and that presynaptic exocytosis is highly effective when the RIM priming activity is artificially tethered to Ca_V_2s^34^. However, several presynaptic properties indicate that active zone organization might be more complex. For example, the coupling distance between Ca_V_2s and releasable vesicles is variable and sometimes up to 100 nm^35–37^, and modelling indicates that the exocytic reaction is best explained by positioning Ca_V_2s in the perimeter of release sites or by excluding them from these sites^38,39^. Furthermore, some studies detect clusters of single active zone proteins that are separated from one another^11,40,41^, suggesting that active zone assembly is organized via self-clustering of individual proteins rather than via complexes between various proteins. Altogether, it has remained challenging to define how the multiple active zone functions are related to the suborganization of its molecular components.

Liprin-α proteins might be important organizers of active zone assembly given their roles upstream of most components^22–25,41^. They interact with RIM and ELKS^42,43^, and with LAR-type receptor protein tyrosine phosphatases (LAR-RPTPs)^44–46^. Because LAR-PRTPs are transmembrane proteins, this provides for a plausible molecular axis to anchor active zone machineries to the presynaptic plasma membrane. In invertebrates, loss of function of Liprin-α results in loss of presynaptic material^22,23,25^, and LAR-RPTPs likely function in the same pathway^24,47^. Indeed, at the fly neuromuscular junction Liprin-α recruits a specific Munc13 isoform^41^. Liprin-α2 and -α3 also support Munc13 targeting at hippocampal synapses^20,21^. In aggregate, this indicates that the Liprin-α/LAR-RPTP pathway may organize presynaptic compartments.

Here, we show that Ca_V_2 and Munc13 each form nanoclusters that are positioned at variable distances from one another at hippocampal synapses. RIM organizes both Ca_V_2 channels and Munc13, but it executes these functions via independent protein domains and interactions. In heterologous cells, Liprin-α recruits RIM into molecular condensates that co-exist with separate Ca_V_2-organizing complexes. At synapses, quadruple knockout of Liprin-α1 to Liprin-α4 reduces the levels of RIM and Munc13 at the active zone and diminishes synaptic vesicle priming, but Ca_V_2 clustering remains intact. Finally, both in transfected cells and at synapses, PTPσ tethers Liprin-α, RIM and Munc13 into protein complexes, but Ca_V_2 is not co-recruited into these complexes. We conclude that Ca_V_2 clustering and vesicle priming sites are organized independently and that Liprin-α and PTPσ specifically support the assembly of priming machinery. Overall, our work argues against a single-complex model of the active zone. Instead, we identify at least two distinct machineries that have independent assembly mechanisms.

## Results

### Variable segregation of Munc13 and Ca_V_2 nanoclusters

To dissect active zone organization, we first determined the subsynaptic localization of two of its essential components: Munc13-1, a protein that defines release sites by priming synaptic vesicles^6,7,48,49^, and Ca_V_2.1, a voltage gated Ca^2+^ channel critical at most central synapses^50–52^. Using antibody staining, both Munc13-1 and Ca_V_2.1 displayed prominent synaptic signals (Fig. 1a). To determine their relative nanoscale distribution, we used DNA Exchange-PAINT^53–55^. En-face synapses containing both proteins were selected, and auto-correlation functions were used to assess the distributions of Munc13-1 and Ca_V_2.1 within single synapses. These first analyses revealed that local densities were heterogeneous (Fig. 1b-d), which is indicative of nanoclustering^56,57^. We then detected nanoclusters for each protein via density-based spatial clustering analyses. Munc13-1 nanoclusters were present at higher density than Ca_V_2.1 nanoclusters and were smaller (Fig. 1e+f), with estimated cluster radii of ∼23 nm for Munc13-1 and ∼30 nm for Ca_V_2.1.

**Figure. 1.**
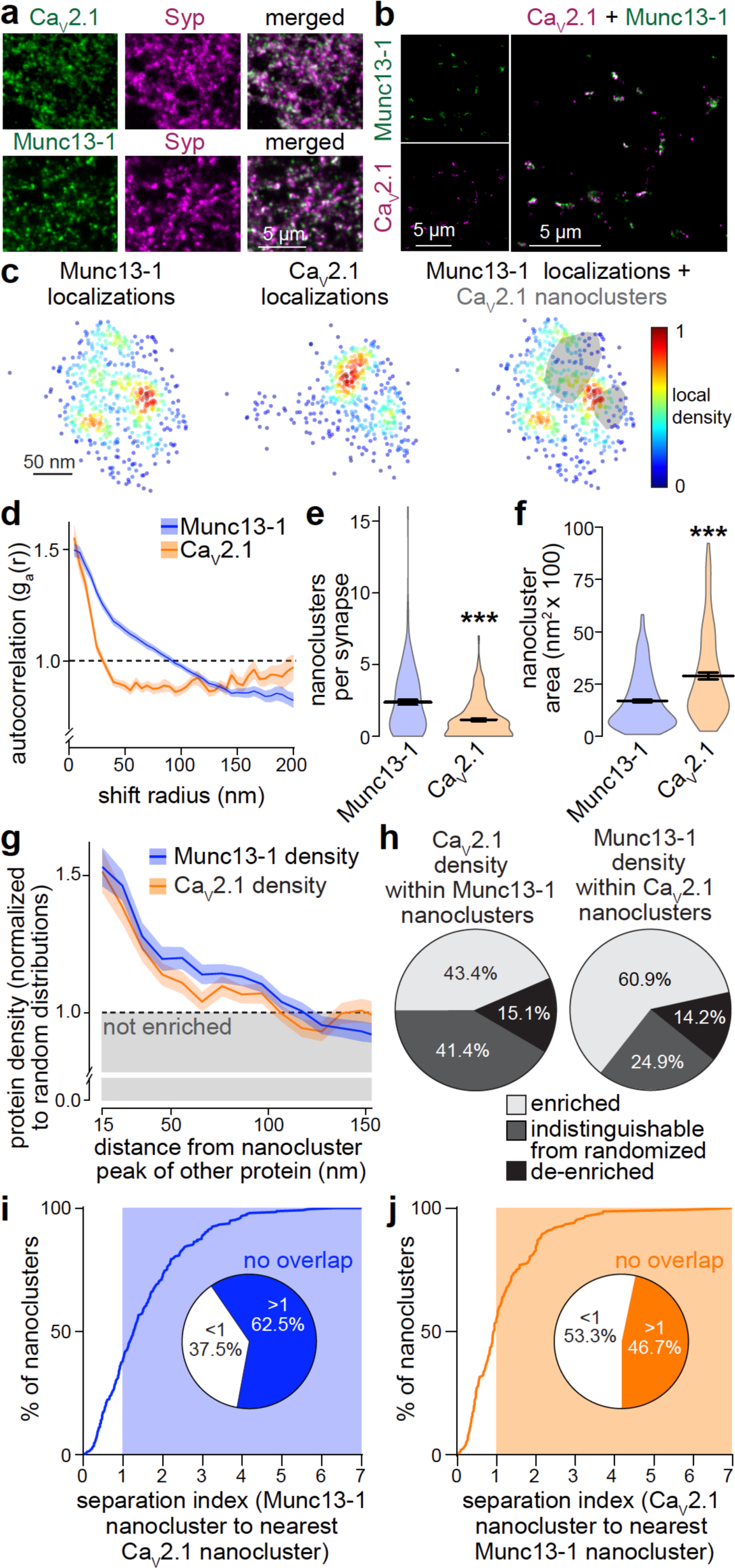
Analyses of Munc13-1 and Ca_V_2.1 nanoclustering (a) Overview confocal microscopic images of cultured hippocampal neurons stained for Munc13-1 or Ca_V_2.1 and co-stained for Synaptophysin (Syp) to mark synapses. (b) Zoomed-out view of rendered images from a DNA Exchange-PAINT experiment to determine the relative localization of Munc13-1 and Ca_V_2.1. (c) Zoomed-in view of protein localizations in an example en-face synapse imaged with Exchange-PAINT showing Munc13-1 (left), Ca_V_2.1 (middle), and Munc13-1 with shaded Ca_V_2.1 nanoclusters identified with a density-based clustering algorithm. (d) Autocorrelation of Munc13-1 and Ca_V_2.1 synaptic densities, measured as the average probability of detecting localizations for the same protein at increasing distances from any given signal; Munc13-1 147 synapses/3 cultures, Ca_V_2.1 147/3. (e) Quantification of the number of Munc13-1 and Ca_V_2.1 nanoclusters per synapse; nanoclusters were identified using a density-based clustering algorithm; N as in b. (f) Quantification of nanocluster area; Munc13-1 350 nanoclusters/3 cultures, Ca_V_2.1 169/3. (g) Density of Ca_V_2.1 localizations at various distances from the peak of Munc13-1 nanoclusters, and vice versa, normalized to a spatially homogenous distribution; N as in f. (h) Percentage of Munc13-1 nanoclusters (left) enriched with Ca_V_2.1, containing Ca_V_2.1 density indistinguishable from its randomized distribution, or de-enriched of Ca_V_2.1, and vice versa for Ca_V_2.1 nanoclusters (right); N as in f. (**i, j**) Histogram showing the index of separation from Munc13-1 nanoclusters to the nearest Ca_V_2.1 nanocluster (i) and vice versa (j), and pie chart showing the percentage of clusters that show any overlap (index <1) or no overlap (index >1); N as in f. Data are mean ± SEM; **p < 0.01, ***p < 0.001 as determined by Mann-Whitney U tests (d, e).

Given these differences in clustering, we hypothesized that these proteins are not strictly colocalized and measured their enrichment relative to one another. We first assessed the density of one protein relative to nanoclusters of the other. Ca_V_2.1 density was high within the average radius of Munc13-1 nanoclusters, and vice versa (Fig. 1g), indicating some colocalization. However, a robust fraction of each protein was enriched at distances beyond the nanoclusters of the other protein. Overall, 43% of Munc13-1 nanoclusters were enriched with Ca_V_2.1, and 61% of Ca_V_2.1 nanoclusters with Munc13-1 (Fig. 1h). For each protein, roughly half of the nanoclusters either had a density of the other protein indistinguishable from a randomized distribution or were even de-enriched of it, suggesting independent clustering mechanisms.

Finally, to assess how nanoclusters of these two proteins are positioned relative to one another, we calculated a “separation index”. This index reveals nanocluster spatial relationships; it is <1 when a nanocluster of one protein overlaps with the closest nanocluster of the other, and >1 when there is no overlap. Consistent with the enrichment measurements, 63% of Munc13-1 nanoclusters did not overlap with any Ca_V_2.1 nanoclusters (Fig. 1i) and 47% of Ca_V_2.1 nanoclusters showed no overlap with those of Munc13-1 (Fig. 1j). We conclude that Munc13-1 and Ca_V_2.1 have partially distinct distributions at the active zone, arguing against the model that these proteins are strictly co-organized into a single complex.

### Independent assembly of Ca_V_2 clustering and priming machineries

The differences in the organization of Ca_V_2 and Munc13 suggest separable pathways for the assembly of the underlying machineries. To assess this hypothesis, we studied whether assemblies for synaptic vesicle priming can be generated without recruiting Ca_V_2 and vice versa, or whether these two pathways are inseparable.

Simultaneous deletion of RIM and ELKS (that is RIM1αβ, RIM2αβγ, ELKS1α and ELKS2α) results in active zone disassembly, loss of most Munc13 and impaired Ca_V_2 clustering^26,34^. We used this active zone disruption to test whether recruitment of Munc13-1 and Ca_V_2.1 clustering is independent, or whether activating a pathway for Ca_V_2.1 clustering restores Munc13-1 recruitment and vice versa. We cultured hippocampal neurons from RIM+ELKS conditional knockout mice and used lentiviral expression of Cre or an inactive, truncated Cre to generate quadruple RIM+ELKS knockout (cQKO^R+E^) or control (control^R+E^) neurons, respectively (Fig. 2a). In cQKO^R+E^ neurons, we used lentiviral expression of either RIM1α (to restore both functions^34^) or of one of the two following complementary RIM fragments. The first, PDZ-C2A-P, consists of the PDZ domain that binds to Ca_V_2, three PxxP motifs (P) that interact with RIM-BP to cluster Ca_V_2s, and the C2A domain^8–11,58^. The second, Zn-C2B, contains the zinc finger that binds to Munc13 and Rab3^4,59,60^ and the C2B domain that binds to Liprin-α and PIP2^43,61^. Each construct also contained endogenous linkers and an HA-tag for detection.

**Figure 2.**
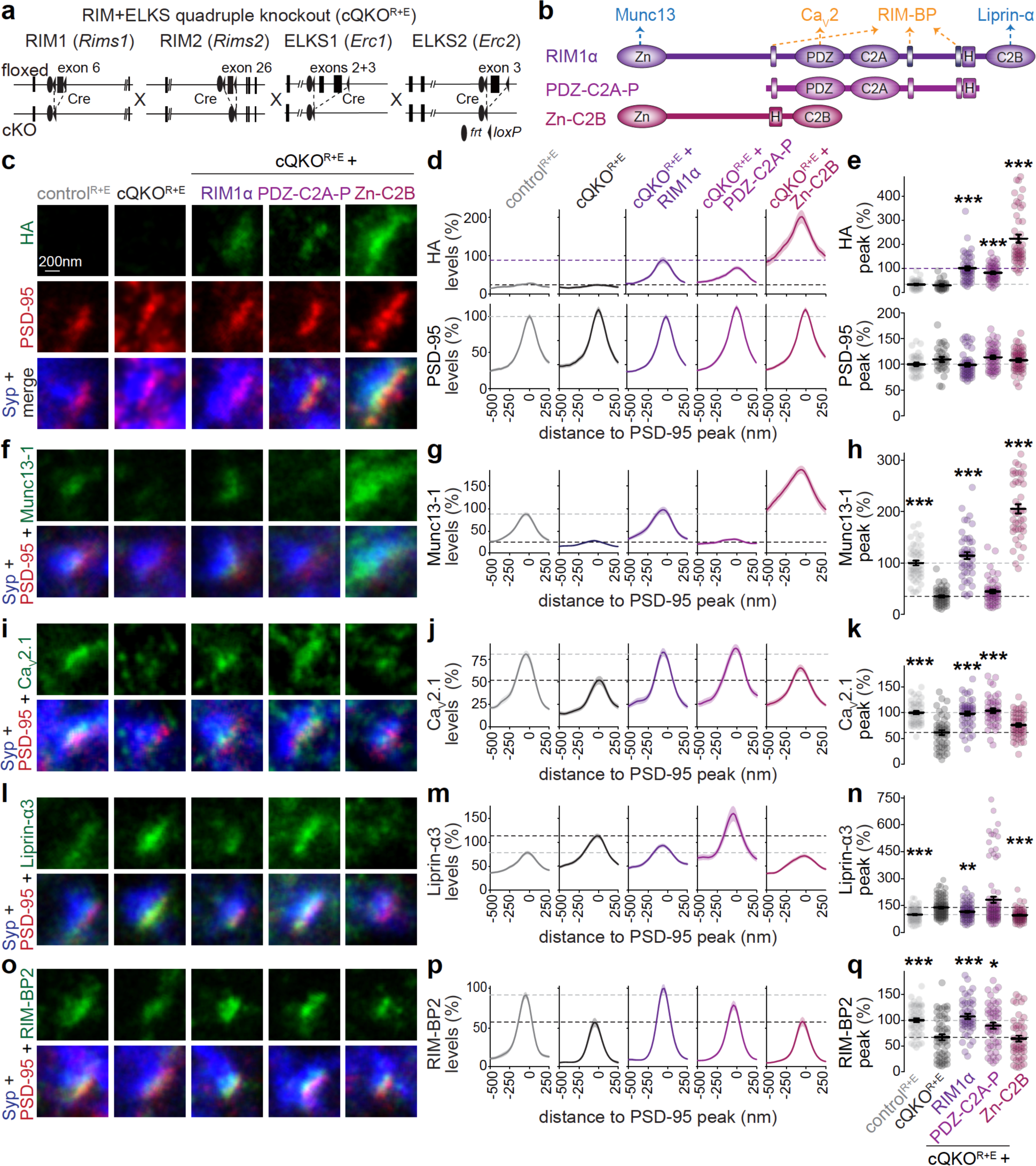
**Distinct RIM domains mediate Ca_V_2 clustering and priming machinery assembly** (**a, b**) Schematics of the *Rims1*, *Rims2, Erc1* and *Erc2* mutant alleles that constitute the conditional RIM+ELKS knockout line (a) and map of rescue constructs (b). Cultured hippocampal neurons after knockout of RIM and ELKS (cQKO^R+E^; lentiviral expression of Cre) or corresponding control neurons (control^R+E^; lentiviral expression of recombination-deficient Cre) were analyzed without or with expression of rescue constructs. HA-tagged RIM1α with key interactions indicated, and PDZ-C2A-P and Zn-C2B (H: HA-tag, Zn: zinc finger domain, PDZ: PDZ domain, C2A and C2B: C2 domains, P: proline rich PxxP-motif) are shown in b. (**c-q**) Example STED images, average line profiles and quantification of the peak intensities of HA and PSD-95 (c-e), Munc13-1 (f-h), Ca_V_2.1 (i-k), Liprin-α3 (l-n) and RIM-BP2 (o-q) at excitatory side-view synapses identified by Synaptophysin (Syp) and PSD-95. Neurons were stained for a protein of interest (HA for RIM1α, Munc13-1, Ca_V_2.1, Liprin-α3 or RIM-BP2; imaged in STED), PSD-95 (imaged in STED), and Synaptophysin (imaged in confocal). A line profile (750 nm x 250 nm) was positioned perpendicular to the center of the elongated PSD-95 object, and synapses were aligned to each other via the PSD-95 peak in the line profile plots (d, g, j, m, p). The maximum value of each profile was used to calculate the peak (e, h, k, n, q). Dotted lines mark levels of cQKO^R+E^ (black), and control^R+E^ (gray) or cQKO^R+E^ + RIM1α (purple). Line profiles and peak intensities were normalized to the average signal in control^R+E^ or cQKO^R+E^ + RIM1α (for HA) per culture; c-e, control^R+E^ 36 synapses/3 independent cultures, cQKO^R+E^ 29/3, cQKO^R+E^ + RIM1α 47/3, cQKO^R+E^ + PDZ-C2A-P 47/3, cQKO^R+E^ + Zn-C2B 48/3; f-h, control^R+E^ 48/3, cQKO^R+E^ 44/3, cQKO^R+E^ + RIM1α 45/3, cQKO^R+E^ + PDZ-C2A-P 49/3, cQKO^R+E^ + Zn-C2B 43/3; i-k, control^R+E^ 45/3, cQKO^R+E^ 47/3, cQKO^R+E^ + RIM1α 45/3, cQKO^R+E^ + PDZ-C2A-P 44/3, cQKO^R+E^ + Zn-C2B 46/3; l-n, control^R+E^ 92/6, cQKO^R+E^ 93/6, cQKO^R+E^ + RIM1α 87/6, cQKO^R+E^ + PDZ-C2A-P 92/6, cQKO^R+E^ + Zn-C2B 72/6; o-q, control^R+E^ 50/3, cQKO^R+E^ 51/3, cQKO^R+E^ + RIM1α 47/3, cQKO^R+E^ + PDZ-C2A-P 51/3, cQKO^R+E^ + Zn-C2B 47/3. Data are mean ± SEM; *p < 0.05, **p < 0.01, ***p < 0.001 shown compared to cQKO^R+E^ as determined by Kruskal-Wallis followed by Holm multiple comparisons post hoc tests (e, h, n, and q), or by a one-way ANOVA followed by Tukey-Kramer multiple comparisons post hoc tests (k). For expression of RIM constructs, confocal microscopic analyses, and STED analyses of inhibitory synapses, see Supplemental figs. 1 and 2.

To assess active zone organization, we used antibody staining followed by sequential confocal and STED imaging with a workflow we established before^20,34,50,62^. For STED analyses, side-view synapses were identified by a synaptic vesicle cloud marked with Synaptophysin and the presence of an elongated area marked with the postsynaptic density protein PSD-95 at the edge of the vesicle cloud. Line profiles perpendicular to the PSD-95 signal were then assessed to evaluate localization and levels of the tested proteins. RIM1 fragments were localized apposed to PSD-95 (Fig. 2c-e) at levels similar to (PDZ-C2A-P) or above (Zn-C2B) those for RIM1α, indicating targeting to the active zone. As described before^26,34,63^, the active zone was disrupted in cQKO^R+E^ neurons with very low levels of Munc13-1, and robust reductions in Ca_V_2.1 and RIM-BP2. The postsynaptic marker PSD-95 was unaltered, and Liprin-α3 was increased by ∼30%, in agreement with previous work^34^ and with an upstream function^22,23,25^.

RIM1α expression in cQKO^R+E^ neurons restored the levels of all measured proteins to those in control^R+E^ synapses. While PDZ-C2A-P restored Ca_V_2.1 and RIM-BP2, it failed to recover Munc13-1. Conversely, Zn-C2B restored Munc13-1 and Liprin-α3, but not Ca_V_2.1 or RIM-BP2 (Fig. 2f-q). Similar results were obtained when we quantified confocal images, and these analyses also established that overall excitatory and inhibitory synapse numbers were not affected (Supplemental fig. 1). STED analyses of inhibitory synapses also matched with these outcomes (Supplemental fig. 2). We conclude that Munc13-containing priming complexes can be assembled without recruiting Ca_V_2 and vice versa, which supports that two independent pathways can mediate the assembly of these presynaptic machineries.

### Liprin-α condensates and Ca_V_2-organizing complexes compete for RIM

If independent assembly pathways for synaptic vesicle priming and Ca_V_2 clustering machineries exist, it should be possible to reconstitute and assess these pathways in a reduced environment. We transfected HEK293T cells with plasmids expressing components of these machineries (Fig. 3a). We previously described that Liprin-α3 forms condensates with RIM1α and Munc13-1 through liquid-liquid phase separation^20^ and that Ca_V_2 complexes can be assembled^10^ in these cells. We first transfected fluorophore tagged Ca_V_2.1, RIM1α and RIM- BP2. This resulted in the formation of protein condensates that contain these three proteins (Fig. 3b), similar to condensates of purified proteins^8,33,64^. These condensates reflect the complexes that cluster Ca_V_2s at the active zone^9–12^. We reasoned that, if only one assembly pathway exists, adding additional active zone proteins should lead to co-condensation with these complexes. Instead, when Liprin-α3 was added (+ Liprin-α3), a fraction of RIM1α was removed from Ca_V_2.1- and RIM-BP2-containing condensates (Fig. 3b+c). Addition of a protein that has no known interactions with RIM, Dynamin-1 (+ unrelated protein), did not have the same effect. We repeated this experiment with tagged Liprin-α3 instead of tagged RIM-BP2 and obtained similar results (Fig. 3d+e). We also used correlative light-electron microscopy (CLEM) in HEK293T cells to assess condensates containing these proteins (Fig. 3f+g). RIM1α and Liprin-α3 localized to electron-dense condensates not surrounded by lipid bilayers (Fig. 3g), similar to the structures we characterized before containing RIM1α, Munc13-1 and Liprin-α^20^.

**Figure 3.**
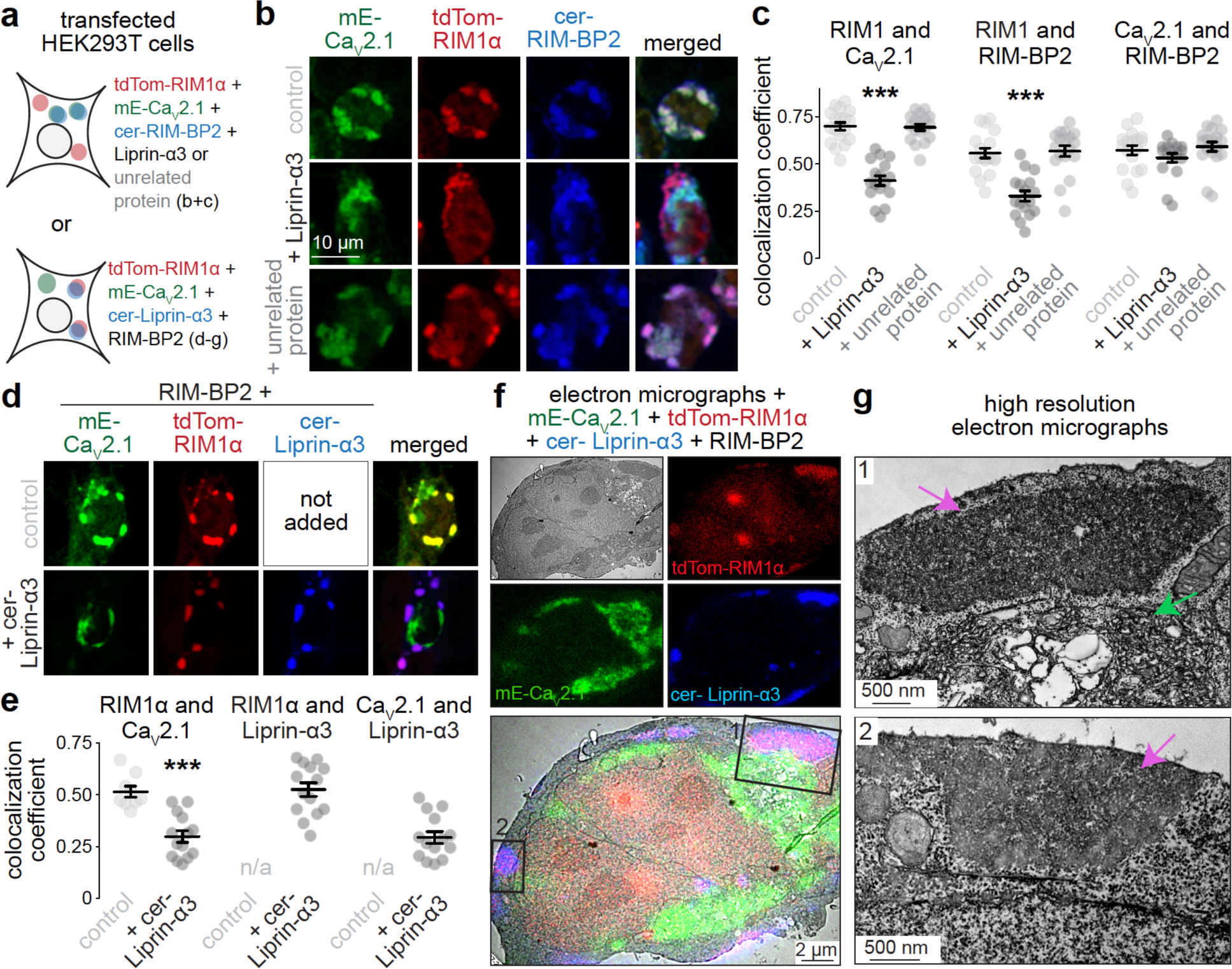
Liprin-α competes for RIM with Ca_V_2.1-organizing complexes (**a**) Overview of the experiment in transfected HEK293T cells. (**b, c**) Example confocal images (b) and quantification (c) of HEK293T cells transfected with cerulean-RIM-BP2 (cer-RIM-BP2), tdTomato-RIM1α (tdTom-RIM1α) and mEOS-Ca_V_2.1 (mE- Ca_V_2.1), and without (control) or with Liprin-α3 (+ Liprin-α3) or Dynamin-1 (+ unrelated protein). The Pearson’s colocalization coefficient is shown (c). In mE-Ca_V_2.1, mEOS was used as a conventional fluorophore for visualization; cer-RIM-BP2 was imaged last to prevent photoconversion of mEOS before other fluorophores were acquired; control 18 images/4 independent transfections, + Liprin-α3 17/4, + unrelated protein 18/4; each image is 250 µm x 250 µm and contains multiple cells. (**d, e**) Example confocal images (d) and quantification (e) of HEK293T cells transfected with untagged RIM-BP2, tdTom-RIM1α and mE-Ca_V_2.1, and without (control) or with cerulean-Liprin- α3 (+ cer-Liprin-α3). The Pearson’s colocalization coefficient is shown (e). Cer-Liprin-α3 was imaged last to prevent photoconversion of mEOS before other fluorophores were acquired, control 9/3, + cer-Liprin-α3 14/3. (**f, g**) Correlative light-electron microscopy (CLEM) example images (f) of HEK293T cells transfected with mE-Ca_V_2.1, tdTom-RIM1α, cer-Liprin-α3 and RIM-BP2, and high resolution zoomed-in images (g) pointing at protein condensates with cer-Liprin-α3 and tdTom-RIM1α (purple arrows) or mE-Ca_V_2.1 (green arrows). Cer-Liprin-α3 was imaged last to prevent photoconversion of mEOS before other fluorophores were acquired; an example cell is shown of a total of 7 cells/2 transfections. Data are mean ± SEM; ***p < 0.001 compared to control as determined by Kruskal-Wallis followed by Holm multiple comparisons post hoc tests (c), or by Mann-Whitney U tests (e).

Ca_V_2.1 was localized distinctly; it was present in the same cell, but in separate electron dense structures that contained membranous organelles (Fig. 3g, zoom-in 1) and it was mostly intracellular. Together with previous work^20,64^, these experiments support that multiple distinct active zone complexes, akin to Ca_V_2 clustering machineries and priming complexes, can be assembled and compete for RIM1α.

### Quadruple knockout of Liprin-α1 to -α4 disrupts priming but not Ca_V_2 clustering machinery

Liprin-α forms condensates with RIM and Munc13^20^ and competes for RIM with Ca_V_2-organzing complexes (Fig. 3). Hence, Liprin-α might organize priming machinery, but not Ca_V_2 clustering at synapses, supporting the model of distinct complexes. We tested this hypothesis through generation and characterization of knockout mice to simultaneously remove all four Liprin-α proteins (Liprin-α1 to -α4) encoded by four genes (*Ppfia1-4*)^65^. Conditional Liprin-α1 (Liprin-α1^f/f^) and Liprin-α4 (Liprin-α4^f/f^) knockout mice were generated by homologous recombination (Supplemental fig. 3). They were then crossed to previously generated conditional Liprin-α2 (Liprin-α2^f/f^)^20^ and constitutive Liprin-α3 (Liprin-α3^-/-^)^21^ knockout mice to generate a quadruple homozygous line (Fig. 4a). We cultured hippocampal neurons from these mice and infected them with lentivirus expressing Cre to generate cQKO^L1-L4^ neurons. We chose day in vitro 7 for viral transduction to avoid developmental effects of Liprin-α deletion that are accompanied by impaired neuronal health. Infection with a virus expressing inactive, truncated Cre and a virus expressing HA-tagged Liprin-α3 was used to generate control^L1-L4^ neurons (Fig. 4a+b). We established previously that lentiviral expression of Liprin-α3 in constitutive Liprin-α3 knockout neurons results in normal Liprin-α3 active zone levels and we did not observe gain-of-function effects^20,21^.

**Figure 4.**
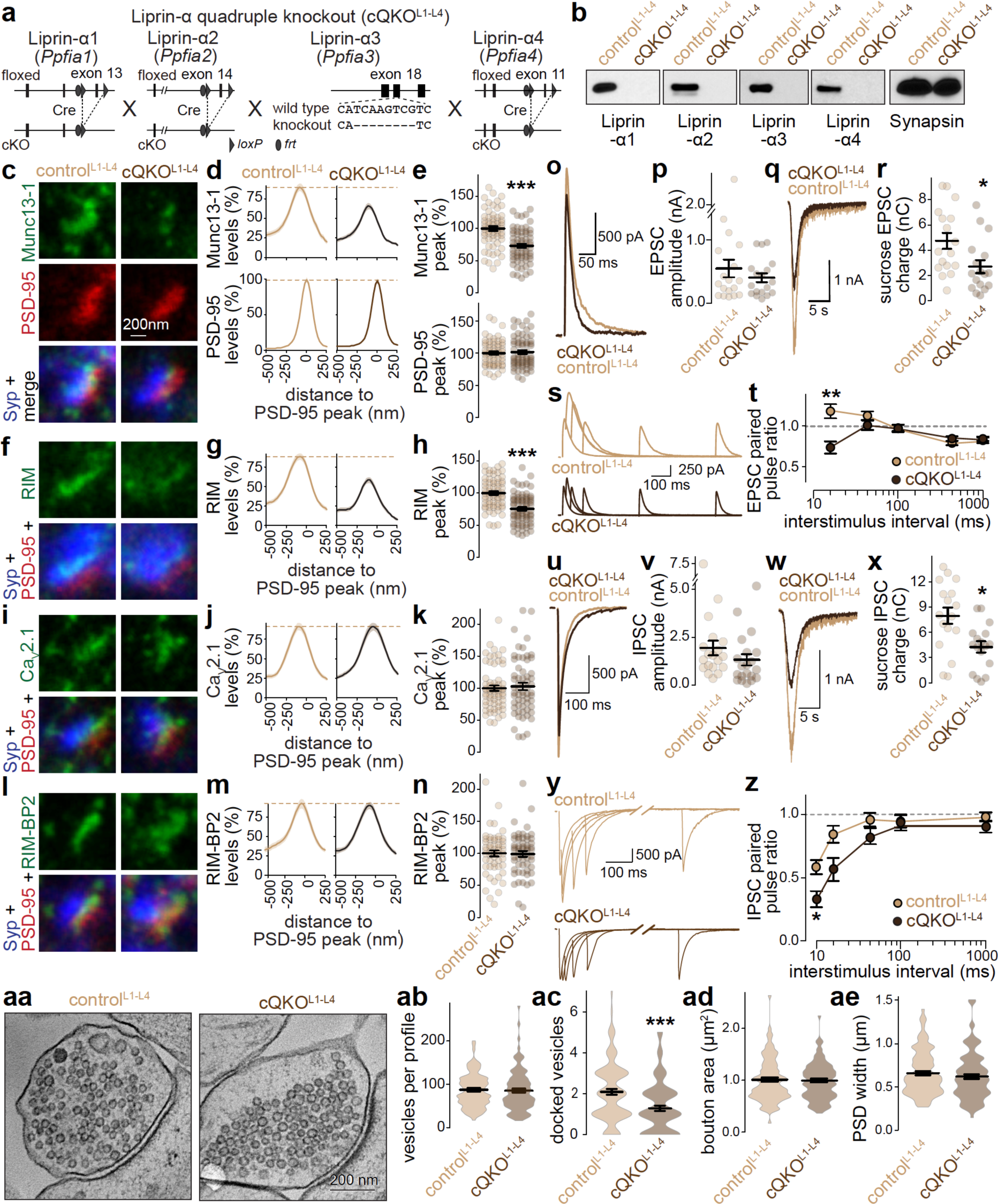
Ablation of Liprin-α disrupts priming machinery but not Ca_V_2 clusters (**a**) Schematics of the *Ppfia1 to 4* alleles that constitute the Liprin-α quadruple mutant mice. These mice were used to generate cultured hippocampal neurons with quadruple knockout of Liprin-α1 to -α4 (cQKO^L1-L4^; lentiviral expression of Cre) or corresponding control neurons (control^L1-L4^; lentiviral expression of recombination-deficient Cre and co-expression of Liprin-α3). (**b**) Western blot to evaluate Liprin-α expression in control^L1-L4^ and cQKO^L1-L4^ neurons. (**c-n**) Example STED images, average line profiles and quantification of the peak intensities of Munc13-1 and PSD-95 (c-e), RIM (f-h), Ca_V_2.1 (i-k) and RIM-BP2 (l-n) at excitatory side-view synapses identified by Synaptophysin (Syp) and PSD-95. Dotted lines in line profile plots mark the levels in control^L1-L4^ neurons; line profiles and peak intensities were normalized to the average signal in control^L1-L4^ per culture; c-e, control^L1-L4^ 62 synapses/3 independent cultures, cQKO^L1-L4^ 65/3; f-h, control^L1-L4^ 65/3, cQKO^L1-L4^ 71/3; i-k, control^L1-L4^ 63/3, cQKO^L1-L4^ 61/3; l-n, control^L1-L4^ 54/3, cQKO^L1-L4^ 56/3. (**o, p**) Example traces (o) and average amplitudes (p) of single action potential-evoked NMDAR- mediated EPSCs; control^L1-L4^ 18 cells/3 independent cultures, cQKO^L1-L4^ 18/3. (**q, r**) Example traces (q) and average AMPAR-mediated charge (r) in response to superfusion with 500 mOsm sucrose; 17/3 each. (**s, t**) Example traces (s) and average NMDA-EPSC paired pulse ratios (t) at increasing interstimulus intervals; N as in o, p. (**u-z**) Same as o-w, but for IPSCs; u+v, control^L1-L4^ 19/3, cQKO^L1-L4^ 20/3; w+x, 17/3 each; y+z, 19/3 each. (**aa-ae**) Example electron micrographs of a synapse cross-section (aa) and quantification of the number of synaptic vesicles per cross-section (ab), the number of docked vesicles per active zone (ac), bouton area (ad) and the width of the postsynaptic density (ae); control^L1-L4^ 103 synapses with 105 active zones/2 independent cultures, cQKO^L1-L4^ 104/109/2. Data are mean ± SEM; *p < 0.005, ***p < 0.001 compared to cQKO^L1-L4^ as determined by Student’s t-tests (e, h, n, and x), Mann-Whitney U tests (k, p, r, v, and ab-ae) or two-way ANOVA followed by Dunnett’s multiple comparisons post hoc tests (t and z). For validation of the Liprin-α1 and -α4 mutant alleles, confocal analyses, and STED analyses of inhibitory synapses, see Supplemental figs. 3-5.

We first assessed active zone assembly at cQKO^L1-L4^ synapses using STED and confocal microscopy. At active zones of excitatory synapses, there was a significant reduction of RIM and Munc13-1, but intact levels of Ca_V_2.1, RIM-BP2 and PSD-95 (Fig. 4c-n). This selective reduction was also observed when we quantified confocal microscopic images (Supplemental fig. 4), and similar effects were present at inhibitory synapses with both confocal and STED analyses (Supplemental figs. 4, 5a-l). This suggests that ablation of Liprin-α selectively disrupts the recruitment of priming machinery.

**Figure 5.**
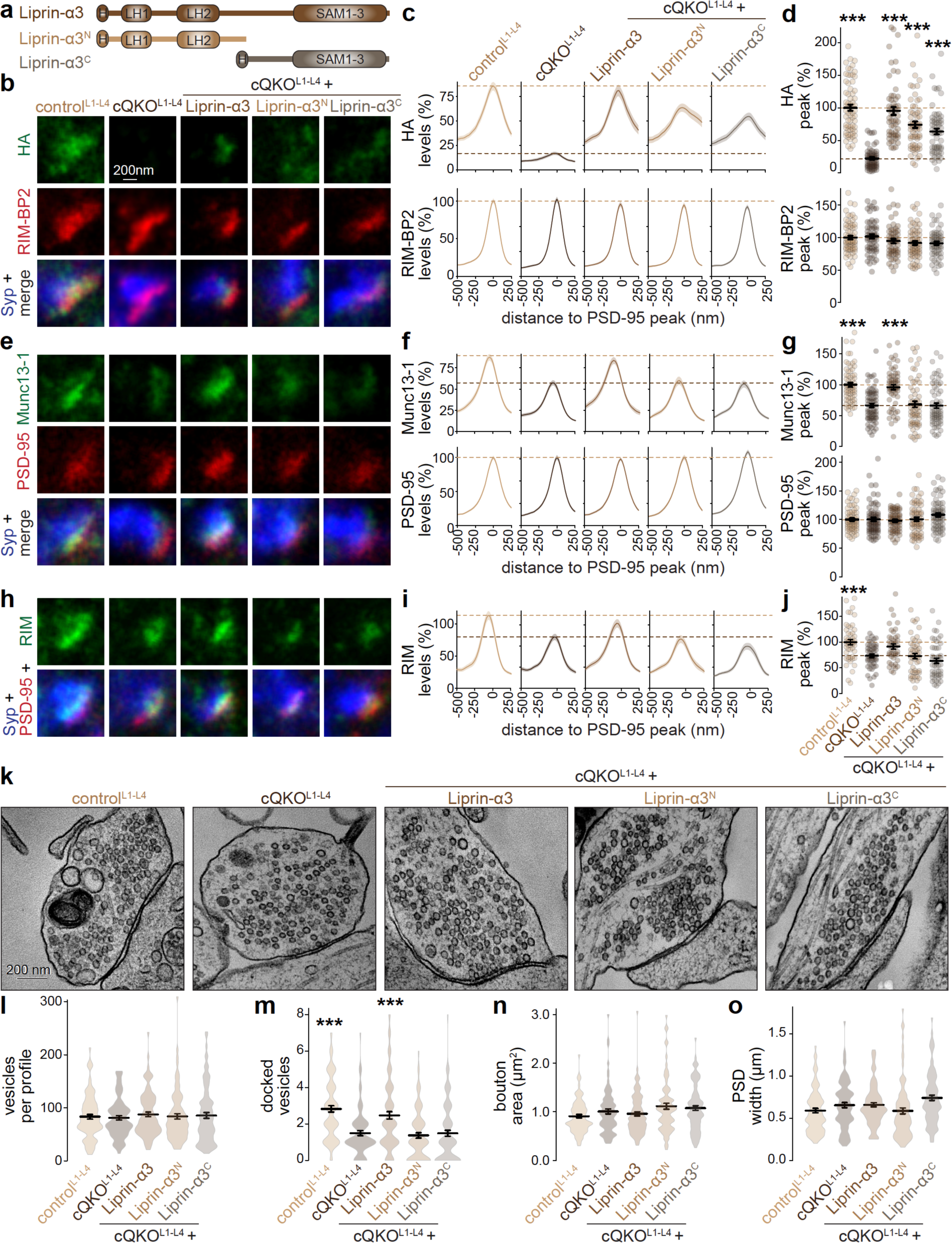
Liprin-α3 operates as a tether via N- and C-terminal domains (**a**) Schematics of HA-tagged Liprin-α3, Liprin-α3^N^, and Liprin-α3^C^ (H: HA-tag, LH: Liprin homology region, SAM: sterile alpha motif). (**b-n**) Example STED images, average line profiles and quantification of the peak intensities of HA and RIM-BP2 (b-d), Munc13-1 and PSD-95 (e-g), and RIM (h-j) at excitatory side-view synapses identified by Synaptohysin (Syp) and PSD-95. Dotted lines mark levels of control^L1-L4^ (light brown) and cQKO^L1-L4^ (dark brown), note that control^L1-L4^ has an HA signal because of the lentiviral expression of HA-tagged Liprin-α3, line profiles and peak intensities were normalized to the average signal in control^L1-L4^ per culture; b-d, control^L1-L4^ 67 synapses/4 independent cultures, cQKO^L1-L4^ 59/4, cQKO^L1-L4^ + Liprin-α3 57/4, cQKO^L1-L4^ + Liprin-α3^N^ 63/4, cQKO^L1-L4^ + Liprin-α3^C^ 60/4; e-g, control^L1-L4^ 65/4, cQKO^L1-L4^ 70/4, cQKO^L1-L4^ + Liprin-α3 60/4, cQKO^L1-L4^ + Liprin-α3^N^ 63/4, cQKO^L1-L4^ + Liprin-α3^C^ 66/4; h-j, control^L1-L4^ 50/3, cQKO^L1-L4^ 47/3, cQKO^L1-L4^ + Liprin-α3 43/3, cQKO^L1-L4^ + Liprin-α3^N^ 49/4, cQKO^L1-L4^ + Liprin-α3^C^ 49/4. (**k-o**) Example electron micrographs of synapse cross-sections (k) and quantification of the number of synaptic vesicles per cross-section (l), the number of docked vesicles per active zone (m), bouton area (n) and the width of the postsynaptic density (o); control^L1-L4^ 91 synapses with 92 active zones/2 independent cultures, cQKO^L1-L4^ 86/87/2, cQKO^L1-L4^ + Liprin-α3 95/96/2, cQKO^L1-L4^ + Liprin-α3^N^ 92/92/2, cQKO^L1-L4^ + Liprin-α3^C^ 96/96/2. Data are mean ± SEM; ***p < 0.001 compared to cQKO^L1-L4^ as determined by Kruskal-Wallis followed by Holm multiple comparisons post hoc tests. For confocal analyses and STED analyses of inhibitory synapses, see Supplemental figs. 6+7.

We next used whole-cell electrophysiological recordings to assess synaptic transmission. The frequency of spontaneous miniature excitatory and inhibitory postsynaptic currents (mEPSCs and mIPSCs, respectively) was decreased in cQKO^L1-L4^ neurons, but their amplitudes were unchanged (Supplemental fig. 5m-t). We evoked synaptic responses with a focal stimulation electrode and recorded resulting EPSCs or IPSCs, isolated pharmacologically in separate batches of cells (Fig. 4o-z). NMDAR-EPSCs were assessed because electrical stimulation induces extensive network activity when AMPARs are not blocked. Overall, evoked EPSCs and IPSCs were similar between genotypes (Fig. 4o+p, 4u+v). Synaptic strength is determined by the size of the readily releasable pool generated by Munc13-mediated vesicle priming, and by vesicular release probability (P)^66,67^. The readily releasable pool, estimated by superfusion with hypertonic sucrose, was reduced by ∼45% at both excitatory and inhibitory cQKO^L1-L4^ synapses (Fig. 4q+r, 4w+x). Changes in P at cQKO^L1-L4^ synapses were assessed by measuring the ratio of two consecutive stimuli at short intervals. Decreases in these ratios at both synapse types indicated an increase in P (Fig. 4s+t, 4x+z). Overall, these results suggest a decrease in the readily releasable pool matching with reductions in RIM and Munc13 (Fig. 4c-h). This is accompanied by an increase in P to account for normal single evoked PSCs.

To complement the functional and STED analyses, we performed electron microscopy on synapses of high-pressure frozen neurons (Fig. 4aa-ae). Matching with the reductions in Munc13 and the readily releasable pool, the number of docked vesicles, identified as synaptic vesicles with no detectable gap between the vesicle and plasma membranes, was reduced in cQKO^L1-L4^ neurons. Total vesicle numbers, bouton area and PSD width were unchanged.

Overall, the analyses of Liprin-α quadruple knockout synapses show that Liprin-α selectively supports the assembly of priming machinery.

### Tethering functions of Liprin-α necessitate simultaneous action of its N- and C-terminal domains

Liprin-α contains two N-terminal Liprin homology regions (LH1 and LH2) that bind to RIM and ELKS and that undergo oligomerization^42,43,64^. These sequences are followed by a linker and three sterile alpha motifs (SAM) that mediate binding to other presynaptic scaffolds including the cell adhesion proteins LAR-RPTPs^44–46^. We hypothesized that specific fragments of Liprin-α might be sufficient to mediate its assembly roles, as observed for RIM (Fig. 2). We generated lentiviruses to express complementary fragments of Liprin-α3 containing either the N-terminal LH sequences (Liprin-α3^N^) or the C-terminal SAM domains (Liprin-α3^C^) and compared their rescue activity to full length Liprin-α3 (Fig. 5a). All constructs were expressed effectively and were targeted to the active zone (Fig. 5b-d, Supplemental fig. 6a-d). Full length Liprin-α3 restored Munc13-1 synaptic and active zone levels at excitatory and inhibitory synapses (Fig. 5e-g, Supplemental figs. 6e-g, 7a-c). This was probably mediated by RIM, whose levels were also restored effectively when measured with confocal microscopy at excitatory and inhibitory synapses and at inhibitory synapses when measured by STED, and showed a strong trend towards rescue at excitatory synapses in STED as well (p = 0.058; Fig. 5h-j, Supplemental figs. 6h-j, 7d-f). Neither Liprin-α3^N^ nor Liprin-α3^C^ produced an increase in the levels of RIM or Munc13-1 (Fig. 5e-j, Supplemental figs. 6e-j, 7). Correspondingly, the number of docked vesicles was only rescued by full length Liprin-α3, but not by the shorter fragments (Fig. 5k-o). These data support that Liprin-α operates as a tether for synaptic vesicle priming machinery and indicate that its function relies on the simultaneous action of N- and C-terminal domains.

### PTPσ anchors priming machinery to membranes in HEK293T cells

RIM and ELKS bind to sequences contained within Liprin-α3’s N-terminus^42,43,64^. Fig. 5 indicates that Liprin-α function requires simultaneous action of the N-terminus and the C-terminus, which binds to LAR-RPTPs^44–46^. We tested whether LAR-RPTPs anchor priming machinery to plasma membranes in transfected HEK293T cells. We used mVenus-tagged Liprin-α3 (mV-Liprin-α3) alone or in combination with one of the three LAR-RPTPs (PTPσ, PTPδ or LAR, not fluorophore-tagged; Fig. 6a). When transfected alone, mV-Liprin-α3 was mostly soluble. We have described before that liquid-liquid phase separation of Liprin-α3 can be induced by activation of PKC in these cells^20^. Co-transfection of PTPσ, PTPδ and LAR promoted Liprin-α3 clustering^45,46^, but to different degrees (Fig. 6b-d). LAR induced less mV-Liprin-α3 condensates than PTPδ (LAR vs. PTPδ, Fig. 6c, p = 0.002) and PTPσ (LAR vs. PTPσ, Fig. 6c, p < 0.001), and these condensates were smaller than those induced by PTPσ (LAR vs. PTPσ, Fig. 6d, p = 0.023). Only mV-Liprin-α3 condensates induced by PTPσ or PTPδ, not those induced by LAR, showed robust recovery after photobleaching (Fig. 6e-g), suggesting that they are liquid droplets similar to Liprin-α3 condensates induced by PKC phosphorylation^20^. This might be functionally important because the liquid state of Liprin-α supports recruitment of active zone material to synapses^20,23^.

**Figure 6.**
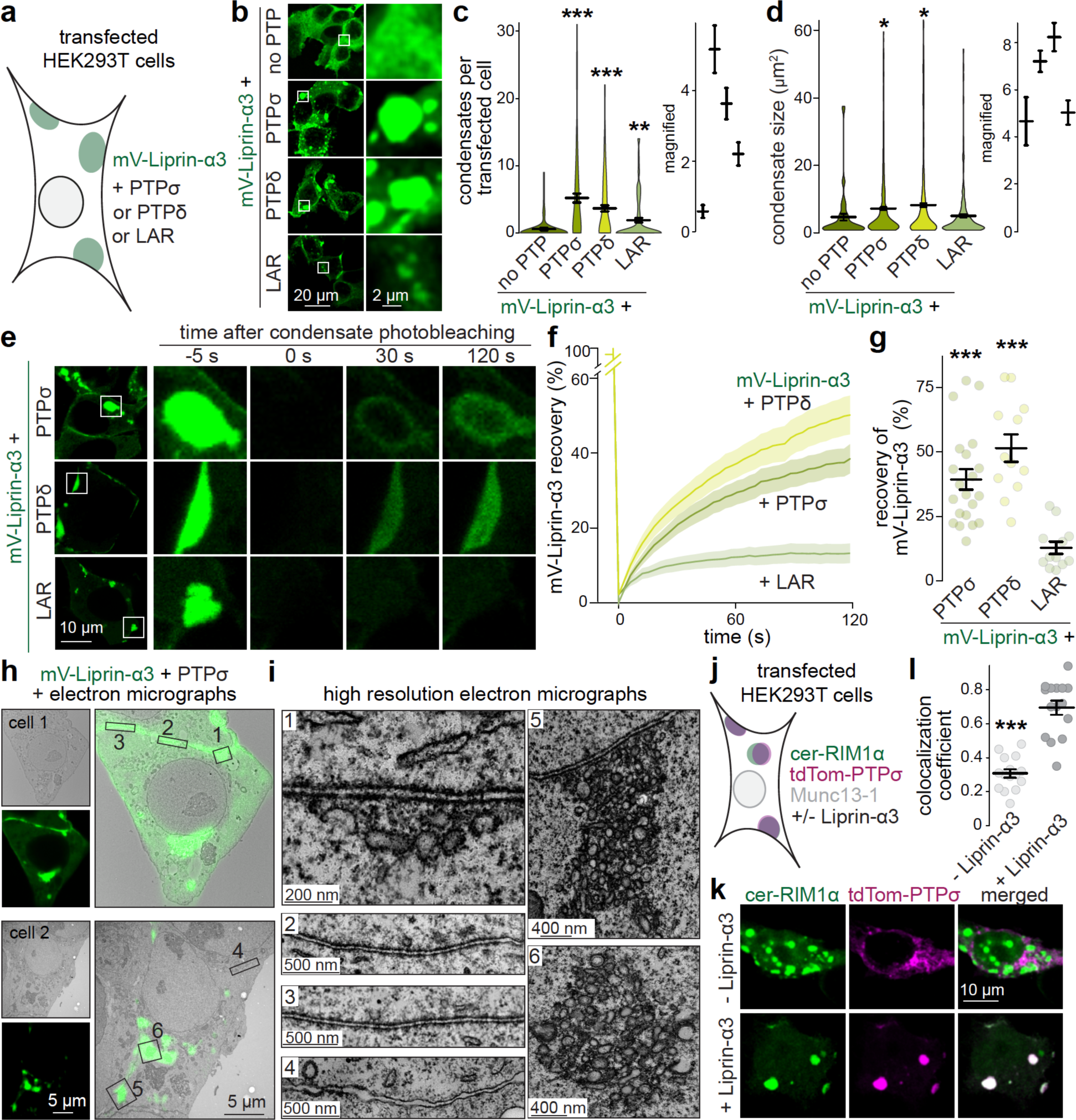
LAR-RPTPs can anchor priming machinery (**a**) Overview of the experiment in HEK293T cells with co-transfection of mVenus-Liprin-α3 (mV- Liprin-α3) and LAR, PTPσ, or PTPẟ (not fluorophore-tagged). (**b-d**) Example confocal images (b) and quantification of the number and size (c+d, with insets on the right showing the relevant range for averages) of mV-Liprin-α3 condensates formed per transfected cell; c, mV-Liprin-α3 + no PTP 91 cells/3 independent transfections, mV-Liprin-α3 + PTPσ 104/3, mV-Liprin-α3 + PTPδ 114/3, mV-Liprin-α3 + LAR 112/3, each image is 250 µm x 250 µm and contains multiple cells; d, mV-Liprin-α3 + no PTP 52 condensates/3 independent transfections, mV-Liprin-α3 + PTPσ 505/3, mV-Liprin-α3 + PTPδ 414/3, mV-Liprin-α3 + LAR 209/3. (**e-g**) Example confocal images (e), time course of recovery after photobleaching (f) and extent of recovery after 120 seconds (g). In e, zoomed-in images were adjusted to visually match fluorescence intensity of condensates before bleaching and adjustments were identical within each condition; mV-Liprin-α3 + PTPσ 21 condensates from independent cells/3 independent transfections, mV-Liprin-α3 + PTPδ 12/3, mV-Liprin-α3 + LAR 12/3. (**h, i**) CLEM example images (h) of two HEK293T cells transfected with mV-Liprin-α3 and PTPσ, and high resolution zoomed-in images (i) of membrane-attached condensates (1-3), membranes without condensates (4) and cytosolic condensates (5+6); two example cells are shown of a total of 5 cells/1 transfection. (**j**) Overview of the experiment in HEK293T cells with co-transfection of cerulean-RIM1α (cer- RIM1α), Munc13-1, and tdTomato-PTPσ (tdTom-PTPσ), and either with or without Liprin-α3. (**k, l**) Example confocal images (k) and quantification of Pearson’s colocalization coefficient of tdTom-PTPσ and cer-RIM1α. The distribution of cer-RIM1α in the - Liprin-α3 condition was variable with either forming condensates (shown) or being more widespread (not shown); colocalization with PTPσ was low in both cases and cells were included in the quantification regardless of cer-RIM1α distribution; 15 images/3 independent transfections per condition, each image is 250 µm x 250 µm and contains multiple co-transfected cells. Data are mean ± SEM; *p < 0.05, ***p < 0.001 compared to mV-Liprin-α3 + no PTP (c+d) or to mV-Liprin-α3 + LAR (g) as determined by Kruskal-Wallis followed by Holm multiple comparisons post hoc tests (c, d, g), or by Mann-Whitney U tests (l).

Because LAR-RPTPs are transmembrane proteins, the induced Liprin-α3 condensates should be associated with membrane compartments. Using CLEM (Fig. 6h+i), we found that mV-Liprin- α3 condensates were electron dense, but not surrounded by lipid bilayers, consistent with liquid droplets. Many of these structures were directly attached to the plasma membrane. The stretches of plasma membrane that were associated with condensates were separated from the apposed cell with even intercellular spacing (Fig. 6i, zoom-ins 1-3). This organization, in appearance similar to a synaptic cleft, was not present between adjacent cells without fluorescent signals (Fig. 6i, zoom-in 4). There were also fluorescent structures not directly attached to the plasma membrane, and these contained membranous organelles that might be small vesicles or tubulovesicular structures (Fig. 6i, zoom-ins 5+6). The same intracellular membranous compartments were observed in larger plasma membrane-associated condensates (Fig. 6i, zoom-in 1).

Finally, we tested whether PTPσ-induced Liprin-α3 condensates co-recruit priming machinery. We co-transfected cerulean-tagged RIM1α (cer-RIM1α) and Munc13-1, since these two proteins co-condensate in HEK293T cells in the presence of Liprin-α3^20^, and included tdTomato-tagged PTPσ (tdTom-PTPσ) with or without Liprin-α3 (Fig. 6j). In the presence of Liprin-α3, tdTom- PTPσ and cer-RIM1α colocalized into protein condensates, while colocalization was poor in the absence Liprin-α3 (Fig. 6k+l). Overall, we conclude that PTPσ induces liquid-liquid phase condensation of Liprin-α3. This process drives the assembly of cellular compartments that include membranous structures and that can be anchored to the plasma membrane, and RIM can be co-recruited into these condensates.

### PTPσ promotes priming machinery assembly but not Ca_V_2 clustering

We finally investigated whether PTPσ anchors vesicle priming machinery at synapses. Previous work, including ours, reported that the combined deletion of PTPσ, PTPδ and LAR does not produce detectable impairment in active zone structure and function when assessed with STED microscopy and electrophysiology^62,68^. This is likely due to redundant functions with other presynaptic cell adhesion proteins, for example neurexin^69^. Furthermore, the distinct properties of PTPσ, PTPδ and LAR in clustering Liprin-α (Fig. 6) suggest diverging roles for these proteins, complicating the assessment of knockout phenotypes. To bypass this limitation, we forced synapses to operate with a single LAR-RPTP, either PTPσ or LAR, by re-expressing them individually in LAR-RPTP triple knockout neurons (Fig. 7a+b)^62^. We used a lentivirus expressing Cre recombinase to produce LAR-RPTP triple knockout (cTKO^RPTP^) neurons and added a lentivirus to express PTPσ (cTKO^RPTP^ + PTPσ) or another lentivirus to express LAR (cTKO^RPTP^ + LAR). Both PTPσ and LAR were expressed effectively and targeted to synapses in cTKO^RPTP^ neurons (Fig. 7c-e, Supplemental fig. 8a-c). We first measured the levels of active zone proteins with STED and confocal microscopy. The cTKO^RPTP^ + PTPσ neurons showed a ∼50% increase in Liprin-α3, and concurrent enhancement in RIM and Munc13-1 at both excitatory and inhibitory synapses, while Ca_V_2.1 was unchanged (Fig. 7f-q, Supplemental figs. 8d-o, 9a-l). In contrast cTKO^RPTP^ + LAR neurons had only modestly (∼20%) increased Liprin-α3, but no detectable increase in RIM and Munc13-1. This indicates that the priming machinery consisting of Liprin-α, RIM and Munc13 can be specifically recruited to synapses by PTPσ.

**Figure 7.**
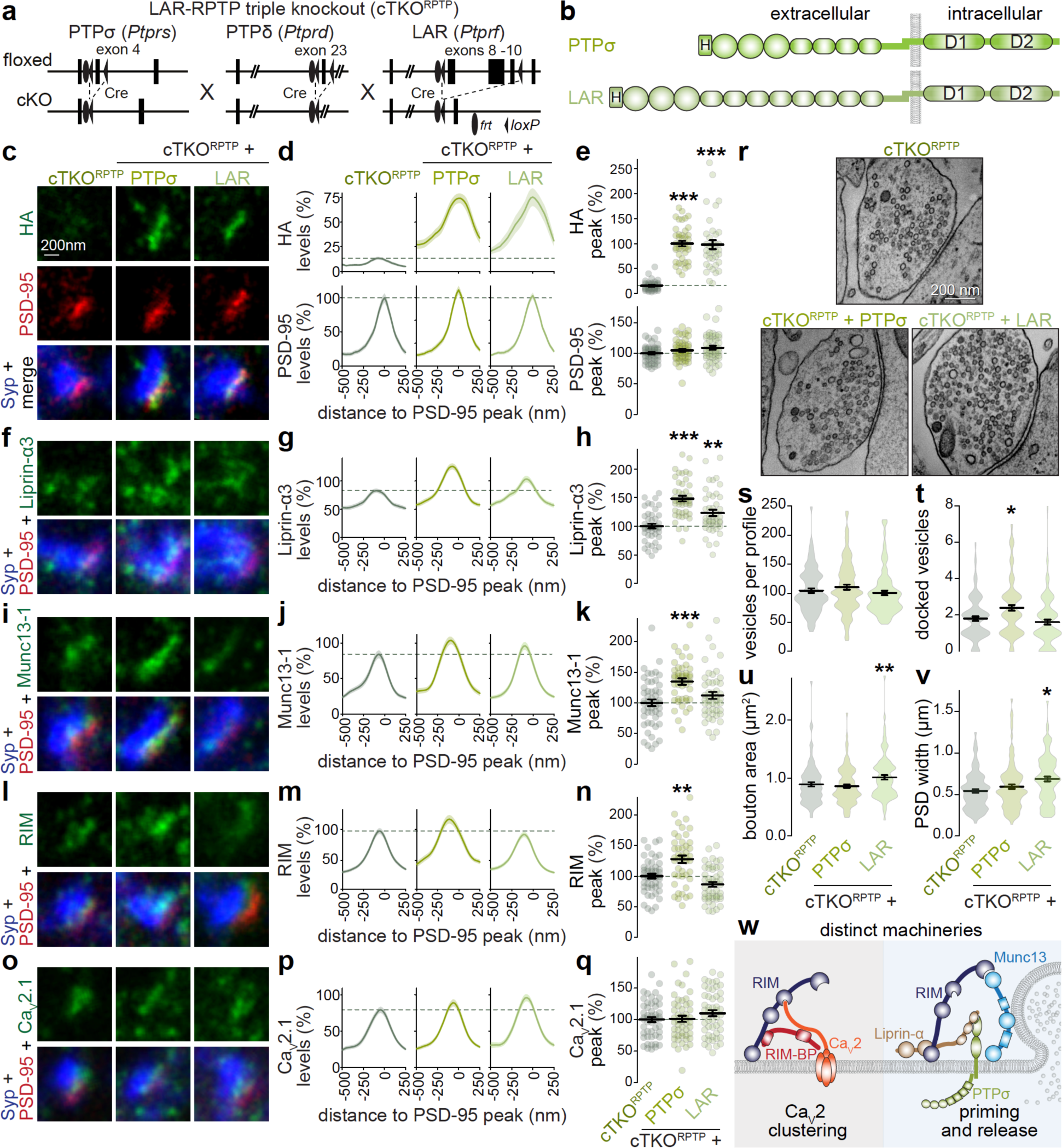
PTPσ assembles priming machinery but does not cluster Ca_V_2s (**a**, **b**) Schematics of the *Ptprs*, *Ptprd* and *Ptprf* mutant alleles that constitute the LAR-RPTP triple conditional knockout mouse line (a) and map of rescue constructs (b). Cultured hippocampal neurons after knockout of PTPσ, PTPδ and LAR (cTKO^RPTP^; through expression of cre-lentiviruses) were analyzed without or with expression of LAR-RPTPs (b). Schematics of HA-tagged PTPσ and LAR are shown (H: HA tag, D1 and D2: phosphatase domains 1 and 2). (**c-q**) Example STED images, average line profiles and quantification of the peak intensity of HA and PSD-95 (c-e), Liprin-α3 (f-h), Munc13-1 (i-k), RIM (l-n) and Ca_V_2.1 (o-q) at excitatory side-view synapses identified by Synaptophysin (Syp) and PSD-95. Dotted lines mark levels of cTKO^RPTP^, line profiles and peak intensities were normalized to the average signal in cTKO^RPTP^ per culture; c-e, cTKO^RPTP^ 38 synapses/3 independent cultures, cTKO^RPTP^ + PTPσ 42/3, cTKO^RPTP^ + LAR 37/3; f-h, cTKO^RPTP^ 45/3, cTKO^RPTP^ + PTPσ 46/3, cTKO^RPTP^ + LAR 45/3; i-k, cTKO^RPTP^ 51/3, cTKO^RPTP^ + PTPσ 45/3, cTKO^RPTP^ + LAR 48/3; l-n, cTKO^RPTP^ 50/3, cTKO^RPTP^ + PTPσ 45/3, cTKO^RPTP^ + LAR 48/3; o-q, cTKO^RPTP^ 51/3, cTKO^RPTP^ + PTPσ 50/3, cTKO^RPTP^ + LAR 51/3. (**r-v**) Example electron micrographs (r) and quantification of the number of synaptic vesicles (s), docked vesicles (t), bouton area (u) and the width of the postsynaptic density (v); cTKO^RPTP^ 97 synapses with 99 active zones/2 independent cultures, cTKO^RPTP^ + PTPσ 105/112/2, cTKO^RPTP^ + LAR 95/103/2. (**w**) Model of active zone organization with two distinct machineries, one to cluster Ca_V_2 and one to prime synaptic vesicles for fusion. Data are mean ± SEM; *p < 0.05, **p < 0.01, ***p < 0.001 compared to cTKO^RPTP^ as determined by Kruskal-Wallis followed by Holm multiple comparisons post hoc tests (e, h, k, n, and s-v), or by a one-way ANOVA followed by Tukey-Kramer multiple comparisons post hoc tests (q). For confocal analyses and inhibitory synapses assessed by STED, see Supplemental figs. 8+9.

We finally assessed effects of the same manipulations on synapse function and ultrastructure. PTPσ expression induced an increase in the pool released in response to hypertonic sucrose, consistent with the recruitment of priming machinery, while expression of LAR did not induce this change (Supplemental fig. 9m-r). Correspondingly, the number of docked vesicles assessed in high-pressure freezing electron microscopy was increased in cTKO^RPTP^ + PTPσ neurons, while the other quantified parameters were unaffected (Fig. 7r-v). LAR expression induced somewhat enlarged boutons and PSDs, perhaps reflecting LAR’s synaptogenic roles^70–73^, but did not produce an increase in docking. These data support the model that PTPσ anchors priming machinery through recruitment of Liprin-α. LAR does not support these functions in the preparation assessed here.

## Discussion

We dissected assembly mechanisms that underlie the nano-organization of the machinery for neurotransmitter release. We found that the active zone is defined by two distinct sub- assemblies, one for clustering Ca^2+^ channels and one for docking, priming and fusing synaptic vesicles. This conclusion relies on five lines of evidence. First, Ca_V_2.1 and Munc13-1 form nanoclusters with distinct distributions, with ∼half of Munc13-1 nanoclusters not enriched with Ca_V_2.1 and vice versa. Second, each complex is assembled by RIM, but distinct RIM sequence motifs mediate assembly of these complexes independently. Third, in transfected cells, RIM forms condensates with Ca_V_2.1 and RIM-BP2, but co-expression of Liprin-α leads to a redistribution of RIM into separate condensates. Fourth, mouse gene knockout and rescue experiments establish that Liprin-α selectively participates in the assembly of priming complexes, but not in clustering of Ca^2+^ channels. Finally, PTPσ selectively anchors priming machinery to plasma membranes, likely via Liprin-α, both in transfected cells and at synapses. Our work establishes that vesicle priming and Ca_V_2 clustering machineries are independent and that Liprin-α and PTPσ specifically support priming machinery assembly.

Previous work suggested that Ca_V_2 is a core component of the machinery that primes and releases synaptic vesicles, arguing for a single active zone protein complex. RIM binds to both Munc13 and Ca^2+^ channels and might directly link docked vesicles to Ca_V_2s^2,10,74^. Fittingly, vesicles can be recruited to the surface of reconstituted liquid condensates containing RIM, RIM-BP and Ca_V_2^8,33^, and artificially attaching the Munc13-binding domain of RIM to Ca^2+^ channels is highly effective in mediating release^34^. If a single protein complex clusters Ca_V_2s and organizes vesicle priming, Ca_V_2s should be at short, invariable distances from releasable vesicles. In contrast, functional studies and modelling suggest that Ca^2+^ channels are at larger, variable distances from these vesicles^35–39^, arguing for more complex active zone organization.

In line with the functional and modelling studies, we found that separable assembly pathways lead to distinct nano-organization of Ca_V_2 and Munc13 (Fig. 7w). The model of separate machineries for vesicle priming and Ca_V_2 clustering allows for flexible positioning of primed vesicles from Ca^2+^ entry, enabling diverse release properties and regulation. In this model, the many parallel interactions between active zone proteins^1,2^ do not result in a “supercluster” containing all active zone components and docked vesicles. Instead, there can be competition between different machineries for specific protein components (Fig. 3)^64^. This might also explain the increases in release probability after selective disruption of the priming complex (Fig. 4): when one assembly is impaired, the other might work more efficiently because competition for space at the plasma membrane and for components that participate in both complexes decreases.

RIM is a central active zone organizer that participates in both complexes. It has been studied in its role in Ca_V_2 clustering together with RIM-BP^8–12,28,75,76^. Parallel studies discovered that RIM docks vesicles and recruits Munc13 for priming^4,10,59,60,75,77–79^. We find that the two machineries are separate and mechanistically extend this finding to reveal that Liprin-α and PTPσ selectively support priming complex assembly (Figs. 3-7)^20,21,41^. While RIM participates in both complexes, RIM-BP, Liprin-α, and Munc13 primarily associate with one of the two complexes. There may be exceptions to this separation. At hippocampal mossy fiber synapses, for example, RIM-BP has roles in vesicle priming^29^. Another active zone protein, ELKS, supports priming at some synapses and Ca^2+^ entry at others^13,15,16,80^. Ultimately, the composition of any given active zone may not always be strictly defined into two types of complexes. Vesicle docking and priming, the coupling of these vesicles to Ca^2+^ channels, and the vesicular fusion reaction are active processes that are accompanied with movement of vesicles and of active zone material.

Presynaptic composition is also modulated over longer time scales. In line with this, we find heterogeneity in active zone organization with a subset of Munc13 nanoclusters enriched with Ca_V_2 (Fig. 1). The model of liquid-liquid phase separation for active zone assembly^8,20,23,33^ is well suited to support these dynamics and may continuously reshape the sub-complexes described here.

One question that arises from the multi-complex model (Fig. 7w) is how the different active zone protein machineries are tethered to one another. The two complexes must be close enough for Ca^2+^ sensors on primed vesicles to detect Ca^2+^ entry for exocytosis. Different parts of a single RIM molecule could simultaneously associate with separate machineries for vesicle priming and for Ca_V_2 clustering. This is possible because the distance between the RIM zinc finger, which binds to Munc13 and vesicular Rab3, and the RIM PDZ domain, which binds to the C-terminus of Ca_V_2 or of ELKS, can be several tens of nanometers long if the linker between them is unfolded. The Ca_V_2 intracellular C-terminus can bridge a similar distance as well. Other proteins, for example ELKS or RIM-BP, could similarly connect the two machineries. Another possibility is that one molecule can only participate in priming or in Ca_V_2 clustering. This scenario is supported by previous work and by the observation that Liprin-α competes for RIM with Ca_V_2-organizing complexes in HEK293T cells (Fig. 3)^64^. It is also strengthened by mutually exclusive biochemical interactions: Ca_V_2 and ELKS, for example, bind to the same binding pocket of the RIM PDZ domain^10,81^ and the specific binding partner could determine which machinery a given RIM protein associates with.

The relative positioning of active zone protein complexes could also be determined by their interactions with transmembrane proteins. The main candidates are neurexins, LAR-RPTPs, and Ca^2+^ channels themselves^50,62,68,69,82,83^ and interactions of these proteins with extracellular material or other membrane proteins could control the spatial organization. Several recent findings on synaptic cell adhesion are suggestive for this model. Work on neurexins revealed that these proteins modulate synapse properties^82,84^, fitting with a role in positioning of specific complexes within an active zone. Furthermore, knocking out one protein family rarely has strong synapse assembly defects^50,62,68,82^. Instead, there is redundancy, as illustrated by studies on neurexins and LAR-RPTPs. On their own, neurexins shape synapse properties likely through organizing Ca_V_2 complexes^84,85^. LAR-RPTPs specifically support the assembly of priming complexes (Figs. 6+7). Removing both proteins at the same time disrupts both complexes and induces strong synapse assembly defects^69^. We propose that the sub-structuring of the active zone that we describe here accounts for the resilience of synapse and active zone assembly that is often observed in loss-of-function studies^12,26,34,63,69,82,86^.

## Acknowledgments

We thank Claire Qiao, Vanessa Charles and Gillian Handy for technical support, all members of the Kaeser laboratory for insightful discussions, Jaewon Ko, Sandra Schmid, Richard Held, Michael Davidson and Tao Xu for plasmids, Susanne Schoch for antibodies, and Matthijs Verhage and Jurjen Broeke for the SynapseEM MATLAB script. This work was supported by grants from the NIH (R01MH113349 and R01NS083898 to P.S.K., R37MH080046 to T.A.B., K99NS129959 to J.E-M., F31MH117920 to P.A.D.), an Alice and Joseph E. Brooks postdoctoral fellowship (to J.E-M.) and a Stuart H.Q. & Victoria Quan fellowship (to J.W.A.). G.dN. is currently at the Massachusetts Institute of Technology and P.A.D is currently at Johns Hopkins University. We acknowledge the following core facilities at Harvard Medical School: the Neurobiology Imaging Facility, the Nikon Imaging Center, the HMS Electron Microscopy Facility, and the Transgenic Mouse Core at the Dana-Farber/Harvard Cancer Center (supported in part by an NCI Cancer Center Support Grant # P30CA006516).

## Author contributions

Conceptualization, J.E-M. and P.S.K; Methodology: J.E-M., J.W.A., S.R.M., A.R.L., P.A.D. and G.dN.; Investigation: J.E-M., J.W.A., S.R.M., and G.dN.; Formal Analysis: J.E-M., S.R.M., A.R.L., P.A.D., G.dN., T.A.B. and P.S.K.; Writing-Original Draft: J.E-M. and P.S.K.; Writing -Review and Editing: J.E-M., J.W.A., S.R.M., A.R.L., P.A.D., G.dN., T.A.B. and P.S.K.; Supervision: T.A.B, and P.S.K.; Funding Acquisition: J.E-M., T.A.B., and P.S.K.

## Materials and methods

### Mouse lines

The quadruple floxed mice for RIM1 (RRID: IMSR_JAX:015832, targeting the *Rims1* gene to remove RIM1α and RIM1β)^87^, RIM2 (RRID: IMSR_JAX:015833, targeting the *Rims2* gene to remove RIM2α, RIM2β and RIM2γ)^10^, ELKS1α (RRID: IMSR_JAX:015830, targeting the *Erc1* gene to remove ELKS1αA and ELKS1αB)^16^ and ELKS2α (RRID: IMSR_JAX:015831, targeting the *Erc2* gene to remove ELKS2αA and ELKS2αB)^80^ were previously described^26,34^. The triple floxed mice for PTPσ (RRID:IMSR_CMMR:ABCA, targeting the *Ptprs* gene to remove PTPσ), PTPδ (RRID:IMSR_EM:11805, targeting the *Ptprd* gene to remove PTPδ) and LAR (RRID:IMSR_EUMMCR:8210, targeting the *Ptprf* gene to remove LAR) were previously described^62^. The Liprin-α quadruple knockout mice were newly generated. Embryonic stem cells containing the Liprin-α1 (*Ppfia1*) or Liprin-α4 (*Ppfia4*) mutant alleles were obtained from the European Mouse Mutant Cell Repository and we acknowledge the EUCOMM Consortium as the source of this material. The specific cells used were Ppfia1^tm1a^(EUCOMM)^Hmgu^ (clone A02; RRID:IMSR_EUMMCR:25506) and Ppfia4^tm1a^(EUCOMM)^Hmgu^ (clone D06; RRID:IMSR_EUMMCR:3103) in which exon 13 (Liprin-α1) or 16 (Liprin-α4) were flanked by loxP sites. The mutant alleles were generated via homologous recombination in embryonic stem cells by the international knockout mouse consortium. The embryonic stem cells were expanded and injected into C57BL/6N-A/a blastocysts at the Transgenic Mouse Core (Dana-Farber/Harvard Cancer Center, Harvard Medical School) to generate chimeric founders. After germline transmission, the original knockin mice were crossed to mice that express Flp-recombinase under a β-actin promotor^88^ to remove the LacZ/Neomycin cassette to generate the conditional “floxed” alleles. To analyze survival, we compared the obtained offspring ratio from heterozygote matings at postnatal day 21 (P21) to the expected Mendelian genotype distribution. The Liprin-α1 and -α4 floxed mice were crossed to previously generated Liprin-α2 floxed mice and constitutive Liprin-α3 knockout mice^20,21^ to generate the quadruple mutant mice that were maintained as quadruple-homozygotes. Mice were genotyped either in the laboratory or by Transnetyx. For in-lab genotyping, oligonucleotide primers AGCAGAACTTGGGTCTCC and GTGACCACAGGTGTTTGGAG (547 and 360 bp bands for the floxed and wild-type alleles, respectively) were used for Liprin-α1; an additional reaction was used occasionally (with CCCTGTCTCTTACAAGAAACC and GTGACCACAGGTGTTTGGAG, 174 bp band for wild type; no band for floxed). The conditional floxed Liprin-α4 mice were genotyped with oligonucleotide primers GCTATCTCCAGCAGGTAGGAC and CACAGTGCCTGGTGTTCACG (383 and 240 bp bands for the floxed and wild-type alleles, respectively); genotyping reactions for Liprin-α2 and Liprin-α3 were performed as described before^20,21^. All animal experiments were approved by the Harvard University Animal Care and Use Committee.

### Primary neuronal cultures

Primary hippocampal cultures were prepared as described before^20,21,26,34,50,62^. Hippocampi were harvested from newborn pups (P0-P1) and were digested and dissociated. Dissociated cells were plated onto glass coverslips (or MatTek plates, for Exchange-PAINT) in “plating medium” (Mimimum Essential Medium [MEM] supplemented with 0.5% glucose, 0.02% NaHCO3, 0.1 mg/mL transferrin, 10% Fetal Select bovine serum, 2 mM L-glutamine, and 25 mg/mL insulin).

Approximately 24 hours after plating, medium was exchanged with “growth medium” (MEM with 0.5% glucose, 0.02% NaHCO3, 0.1 mg/mL transferrin, 5% Fetal Select bovine serum, 2% B-27 supplement, and 0.5 mM L-glutamine). At 48 to 60 hours after plating, cytosine β-D-arabinofuranoside (AraC) was added at a final concentration of 2-6 µM. The cells were kept in an incubator at 37 °C until day in vitro (DIV) 15 to 17.

### Production of lentiviruses and transduction of primary neurons

Lentiviruses were produced in HEK293T cells maintained in DMEM supplemented with 10% bovine serum and 1% penicillin/streptomycin (HEK medium). HEK293T cells were transfected with the calcium phosphate method with a combination of the packaging plasmids REV, RRE and VSV-G, plus a lentiviral plasmid encoding Cre recombinase (lab plasmid code p009), an inactive, truncated version of Cre (p010), or a protein used for rescue at a molar ratio 1:1:1:1 and with a total amount of ∼4 μg DNA per T25 flask. At 20 to 30 hours after transfection, the medium was changed to “growth medium” and 48 to 60 hours after transfection the supernatant was collected. Viral transductions were done either with freshly-collected viral supernatant or with snap-frozen supernatant stored at -80 °C. Neuronal cultures were transduced at DIV5 (for RIM+ELKS), DIV6 (for LAR-RPTP) or DIV7 (for Liprin-α) with lentiviruses expressing GFP-tagged Cre recombinase (p010) or the inactive version (p011) under a human Synapsin promoter^16,34^, and expression was monitored via the presence of GFP in the nucleus. Only cultures in which no neurons without nuclear green fluorescence were readily detected were used for experiments. In trial experiments, earlier addition of Cre lentivirus in cultures of LAR-RPTP or Liprin-α mutant alleles resulted in neuronal death. Transduction for expression of rescue proteins was done at DIV1 or DIV2 with lentivirus expressing versions of Liprin-α3, RIM1α, LAR or PTPσ under a human Synapsin promotor, or with a lentivirus without an insert in the multiple cloning site as a control (pFSW control; p008, as described^26,34^). pFSW RIM1α-HA (rat, p592) has been previously esablished^34,61^. The numbering used below and the sequence of p592 correspond to Uniprot ID Q9JIR but without the alternatively spliced exons N_83_-W_105_, H_1084_- R_1169_, and A_1207_-T_1378_, and with two variants (H263D and A453P, see NCBI Reference Sequence: NM_052829.3). pFSW RIM1α-HA PDZ-C2A-P (rat, p1050) was produced from p592 and contains amino acids M_509_ through L_1446_. pFSW RIM1α-HA Zn-C2B (rat, p1051) was produced from p592 and contains amino acids M_1_ through S_508_ and S_1380_ through S_1615_. An HA-tag with short exogenous linkers was present between E_1379_ and S_1380_ for RIM1α and PDZ-C2A- P (a position that did not alter RIM1α function or localization in previous studies^34,61^, sequence …E_1379_-EAAAG-YPYDVPDYA-AGAP-S_1380_…), and at S508 for Zn-C2B (sequence …S_508_-YPYDVPDYA-AGAP-S_1380_…). pFSW HA-Liprin-α3 (rat, p526) corresponds in sequence and numbering to NCBI Reference Sequence: NP_001257914.1 with the addition of an N-terminal HA-tag and a short linker (sequence: M-YPYDVPDYA-GAPS-C_3_…C_1192_) and has been described before^20,21^. pFSW HA-Liprin-α3^N^ (p890) was produced from p526 and contains amino acids C_3_ through S_576_. pFSW Liprin-α3^C^ (rat, p891) was produced from p526 and contains amino acids S_576_ through C_1192_. Both p890 and p891 contain an HA-tag at the N-terminus (p890: M- YPYDVPDYA-GAPS-C_3_… S_576_; p891: M-YPYDVPDYA-GAP-S_576_…C_1192_). pFSW HA-PTPσ (human, p846) was cloned from an open reading frame obtained from Jaewon Ko (pCMV HA- PTPσ in a modified pDisplay backbone^89^), and sequence and numbering correspond to GenBank: AAI04813.1. pFSW HA-LAR (human, p1047) was produced from a plasmid obtained from the Harvard Plasmid Repository expressing human LAR, and sequence and numbering correspond to GenBank: AAH48768.1. The N-terminal signal peptide for each was replaced with a mouse IgΚ leader sequence followed by an HA-tag and a short exogenous linker (sequence of p846: METDTLLLWVLLLWVPGSTGD-YPYDVPDYA-GAQPARS-E_30_…T_1501_; p1047: METDTLLLWVLLLWVPGSTGD-YPYDVPDYA-GAQPARS-D_30_…T_1898_).

### Immunofluorescence staining for DNA Exchange-PAINT

Neurons grown on MatTek dishes were fixed in 2% paraformaldehyde (PFA), 4% glucose in phosphate-buffered saline (PBS) at DIV15 to DIV17. They were next permeabilized in PBS supplemented with 100 mM glycine (PBS-G) containing 0.3% Triton X-100 and blocked in PBS- G containing 3% BSA and 0.1% Triton X-100. Five primary antibodies were used: rabbit anti- Munc13-1 (lab antibody code A72, 1:500; RRID: AB_887733), rabbit anti-Ca_V_2.1 (A46, 1:500; RRID: AB_2619841), mouse anti-PSD-95 (A152, 1:500; RRID: AB_10698024), mouse anti-Bassoon (A85, 1:500; RRID: AB_11181058), and guinea pig anti-Bassoon (A67, 1:500; RRID: AB_2290619). The four primary antibodies for DNA-PAINT (A72, A46, A152 and A85) were pre-incubated with their respective DNA-conjugated secondary nanobodies (F2-Anti-Rabbit IgG (S65), F3-Anti-Rabbit IgG (S66), F1-Anti-Mouse kLC (S67), F4-Anti-Mouse kLC (S68); custom-made by Massive Photonics using FAST-PAINT F-series DNA^90^) in PBS at a 1:2.5 molar ratio for 20 minutes in tubes separated from one another. A rabbit Fc fragment was added to the preincubation mixes of Ca_V_2.1 and Munc13.1 at 2 times higher concentration than the primary antibody. After preincubation, all antibodies were mixed and neurons were incubated overnight at 4 °C. Next, goat anti-guinea pig Alexa Fluor 488 (S3, 1:500, RRID: AB_2534117) was added for one hour at room temperature. Finally, the samples were postfixed in 4% PFA diluted in PBS with 4% glucose. Three 5-minute washes with PBS-G were performed between steps.

### Single molecule imaging, processing, and analysis

Images were acquired on a Nikon TI2 inverted microscope equipped with a 100x Apo TIRF oil immersion objective (1.49 NA), 488, 561 & 640 nm excitation lasers, appropriate filters, an Ixon+ 897 EM-CCD camera, and Nikon Elements and MicAO software. Before imaging, 90 nm gold nanoparticles at 1:10 dilution were added as fiducial markers. Imager strands (Massive Photonics; F1-Atto643, F3-Cy3B, F4-Atto643 and F2-Cy3B) were diluted in imaging buffer (containing PBS + 500 mM NaCl + PCA/PCD/Trolox, pH7.4^54^) at a final dilution of 0.25 nM (for Munc13-1 and Ca_V_2.1) or 0.5 nM (for Bassoon and PSD-95). DNA-PAINT videos were acquired for 30,000 frames with 100 ms exposure in two exchange rounds to minimize chromatic aberration. First, PSD-95 (with F1-Atto643) and Munc13-1 (with F3-Cy3B) were imaged. Next, Bassoon (with F4-Atto643) and Ca_V_2.1 (with F2-Cy3B) were imaged. One wash with PBS was applied between rounds. Finally, 100 nm Tetraspeck beads were immobilized on separate coverslips coated with poly-L-lysine and imaged to subsequently generate a transform (t-form). Videos were processed with Picasso https://github.com/jungmannlab/picasso^54^) and custom MATLAB scripts. Target protein DNA-PAINT videos were localized by minimizing a nearest neighbor search, separately for each target (minimum net gradient 9000). Resulting localization files were recombined to one file per exchange round using Picasso’s *join* function, drift corrected with Picasso’s *undrift* function (segment 1000) and corrected for chromatic aberrations using the MATLAB’s *transformPointsInverse* function and a t-form calculated from Tetraspeck bead images (which was calculated using MATLAB’s *fitgeotrans* function with a 2^nd^ degree polynomial, as before^57^). Any residual linear offset between targets was corrected by cross-correlating each other target to the Munc13-1 localizations using a custom MATLAB function. Localizations with standard deviation of the fit <0.3 or >1.6 pixels, localization error >20 nm, or photon count larger than the mode bin of a histogram of photon counts were removed. Finally, localizations were temporally linked using Picasso’s *link* function (0.3 pixel radius, five dark frames allowed). En-face synapses identified by Bassoon and PSD-95 were analyzed. Putative synaptic clusters of Bassoon and PSD-95 were first identified with Picasso’s *dbscan* function (48 nm radius, 10 min points), and clusters with fewer than 75 localizations were removed. Clusters representing nonspecific imager binding were then removed if either their mean frame number was +/- 2 times the standard deviation of a gaussian fit of the mean frame number of all clusters or if the standard deviation of the frame number was <2500 or >11,000, as nonspecific events are localized for only short time and thus their mean frame number deviates from N frames/2 (15,000) and the standard deviation of frame number is small. Only en-face synapses in which the smoothed Bassoon alpha shape was ≤ 2 and the area overlap between Bassoon and PSD-95 alpha shapes was at least 70% were included. Munc13- 1 and Ca_V_2.1 localizations were next restricted to active zone borders marked by Bassoon.

Synapses with <25 localization for Munc13-1, Ca_V_2.1 or PSD-95 within the Bassoon boundaries were excluded. Synapses were analyzed using a custom MATLAB script similar to those previously described, with modifications for 2d localizations^56,57,91^. Autocorrelation was calculated as described^56,57^, modified for 2d localizations. Nanoclusters of Munc13-1 and Ca_V_2.1 were identified using MATLAB’s *dbscan* function. Localizations were randomized 100 times per protein and synapse within Bassoon. Nanoclusters with <5 localizations and those with an area larger than the maximum non-outlier area (area outlier identified using Graphpad Prism’s ROUT method with Q = 0.1%) were excluded. The area of a nanocluster was defined by its alpha shape (with alpha radius 0.9375). Cross-enrichment was calculated as in^56,57^. Specifically, enrichments were normalized to a random distribution and smoothed by replacing high outliers (identified using Prism’s ROUT with Q = 0.1%) with the highest non-outlier value at each distance bin.

To assess enrichment, we first calculated the total enrichment curve from the center of each nanocluster of the opposite protein. Nanocluster enrichment was categorized by comparing the average value of the cross-enrichment with the opposite protein within 60 nm of the nanocluster center with that following 50 randomizations of the opposite protein’s distribution. Nanoclusters were considered enriched or de-enriched when the real average enrichment to the opposite distribution was greater or less than the average plus or minus 1.96 standard deviations of the enrichments to the random distributions, respectively. All other nanoclusters were designated as indistinguishable from randomized. The separation index was defined as SI = d / (r_1_ + r_2_), where *d* is the center-to-center distance of each object and *r_1_* and *r_2_* are the distances between each object’s center and its border along the line that connects the centers of both objects. *d* is calculated as the Euclidean distance between centroids of a nanocluster of interest to the nearest nanocluster of another protein. *r_1_* is calculated as the distance along this line from the centroid to the border of the first nanocluster, and *r_2_* is the same for the second nanocluster.

Example overview images were generated by rendering in FIJI using the ThunderSTORM plugin’s *Averaged Shifted Histogram* method with a magnification 8 (20 nm pixels).

### Western blotting

Lysates from DIV15 to 16 cultures were collected in standard 3x SDS sample buffer diluted with PBS. Brains of P21-28 mice were homogenized with a glass-Teflon homogenizer in a solution containing 150 mM NaCl, 25 mM HEPES, 4 mM EDTA and 1% Triton X-100 (pH 7.5), and then diluted with standard 3x SDS sample buffer. Samples were boiled at 100 °C for 10 minutes and run on SDS-PAGE gels. Proteins were electrophoretically transferred from gels to nitrocellulose membranes for 6.5 h at 4 °C in buffer containing (per L) 200 mL methanol, 14 g glycine and 6 g Tris. Membranes were blocked for 1 hour at room temperature in TBST (Tris-buffered saline with 0.1% v/v Tween-20) containing 10% non-fat milk powder and 5% normal goat serum.

Membranes were incubated with primary and secondary antibodies, and each step was done overnight at 4 °C. The antibody incubation solution contained TBST with 5% milk and 2.5% goat serum. Five 5-minute washes were performed between steps. The primary antibodies used were: rabbit anti-Liprin-α1 (A121, 1:500), rabbit anti-Liprin-α2 (A13, 1:500), and rabbit anti-Liprin-α4 (A2, 1:500) were gifts from S. Schoch^92^; mouse anti-HA (A12, 1:1000; RRID: AB_2565006); mouse anti-Synaptophysin (A100, 1:5000; RRID:AB_887824), mouse anti-Synapsin (A57, 1:5000; RRID: AB_2617071), goat anti-PTPσ (A114, 1:200, RRID:

AB_2607944), and mouse anti-LAR (A156, 1:500, clone E9B9S from Cell signaling). The secondary antibodies were peroxidase-conjugated goat anti-mouse IgG (S52, 1:10000, RRID:AB_2334540), peroxidase-conjugated goat anti-rabbit IgG (S53, 1:10000, RRID:AB_2334589), and peroxidase-conjugated anti-goat IgG (S60, 1:10000).

### Immunofluorescence staining for STED and confocal microscopy

Neurons grown on 12mm, #1.5 glass coverslips were processed as previously described^20,62^. They were fixed at DIV15 to 16 in PBS containing 4% PFA for 10 minutes (for anti-Ca_V_2.1 antibodies 2% PFA were used^50^), followed by blocking and permeabilization in blocking solution (PBS, 3% BSA, 0.1% Triton X-100) for 1 hour at room temperature. Incubations with primary and secondary antibodies were each done overnight at 4 °C in blocking solution. Samples were post-fixed in 4% PFA for 10 min and mounted using ProLong diamond. Three 5-minute washes with PBS were performed between steps. Primary antibodies used were: rabbit anti-Liprin-α3 (A232, 1:500; homemade, knockout-verified for STED^20^), rabbit anti-RIM (A58, 1:500, RRID: AB_887774, knockout-verified for STED^34,61^), rabbit anti-Ca_V_2.1 (A46, 1:500; RRID:

AB_2619841, knockout-verified for STED^50^), rabbit anti-Munc13-1 (A72, 1:500; RRID: AB_887733, knockout-verified for STED^63^), rabbit anti-RIM-BP2 (A126, 1:500; RRID: AB_2619739), mouse anti-HA (A12, 1:500; RRID: AB_2565006), mouse anti-Synaptophysin (A100, 1:500; RRID: AB_887824), mouse anti-PSD-95 (A152, 1:500; RRID: AB_10698024), and mouse anti-Gephyrin (A8, 1:500; RRID: AB_2232546). Secondary antibodies used were: goat anti-rabbit Alexa Fluor 488 (S5; 1:500, RRID: AB_2576217), goat anti-mouse IgG1 Alexa Fluor 488 (S7; 1:500, RRID: AB_2535764), goat anti-mouse IgG1 Alexa Fluor 555 (S19, 1:500, RRID: AB_2535769), goat anti-mouse IgG2a Alexa Fluor 633 (S30, 1:500, RRID: AB_1500826), goat anti-guinea pig Alexa Fluor 405 (S51, 1:500, RRID: AB_2827755).

### Image acquisition and analyses for confocal and STED microscopy

Images were acquired with previously established protocols^20,62^ using a Leica SP8 Confocal/STED 3X microscope equipped with an oil-immersion 100x objective (1.44 NA), a white laser, STED gated detectors, and 592, 660 and 770 nm depletion lasers. For every image, quadruple confocal scans for Synaptophysin, PSD-95, Gephyrin and the protein of interest were followed by triple-color STED scans for PSD-95, Gephyrin and the protein of interest (Synaptophysin was not imaged in STED mode because of depletion laser limitations).

Exceptions to these protocols were: 1. When HA antibodies were used, Gephyrin antibodies could not be included because they are not compatible with our established HA antibodies. 2. For the assessment of Liprin-α3 fragments in Fig. 5b-d and Supplemental fig. 6b-d, RIM-BP2 was used as a marker instead of PSD-95 and Gephyrin. The acquired images were 2048 pixels x 2048 pixels large with a pixel size of 22.7 nm x 22.7 nm, and each image contained hundreds of synapses. Acquisition settings for a given staining and channel were identical for all images within a batch of culture in which all conditions from one experiment were compared. To quantify synapse density in confocal images, individual confocal channels were analyzed with an automatic detection algorithm (available at https://github.com/kaeserlab/3DSIM_Analysis_CL and https://github.com/hmslcl/3D_SIM_analysis_HMS_Kaeser-lab_CL) using Otsu thresholding. The synaptic mask was subsequently used to quantify the fluorescence intensity levels of the protein of interest. Confocal data were normalized to the average in the condition that was used for comparison (which was control, conditional knockout, or conditional knockout infected with a rescue lentivirus as noted in each figure legend) per culture. To quantify STED images, side-view synapses were selected by an experimenter blind to the protein of interest. Side-views were defined as synapses containing a synaptic vesicle cloud of >250 nm from the active zone to the inside of the presynaptic terminal and with a bar-like Gephyrin or PSD-95 structure along its edge. A 750-nm long, 250-nm wide line profile was then drawn perpendicular to the postsynaptic density marker and across its center. After applying a 5-pixel rolled average to the protein of interest and PSD-95 or Gephyrin, line profiles of individual synapses were aligned to the peak of the postsynaptic marker and averaged. The maximum value of the resulting individual line scans was used to calculate the peak intensity. In each culture, line profiles and peak intensities were normalized to the average in the condition that was used for comparison (which was control, conditional knockout, or conditional knockout infected with a rescue lentivirus as noted in each figure legend). In line profile plots, peaks appear below 100% because peak intensities for proteins of interest are not always at the same position. For representative images, a smooth filter was added, brightness and contrast were linearly adjusted, and images were interpolated. Identical adjustments were applied to representative images within an experiment. Quantifications were performed on original images without brightness and contrast adjustments and without background subtraction or resampling. Data were acquired and analyzed by an experimenter blind to genotype.

### Imaging and analyses of protein condensates in transfected HEK293T cells

HEK293T cells were plated on 12 mm, #1.0 or #1.5 glass coverslips (for fixed samples) or on MatTek dishes (for fluorescence recovery after photobleaching [FRAP] experiments) in HEK medium containing Dulbecco’s Modified Eagle Medium (DMEM) with 10% Fetal bovine serum (Atlas Biologicals F-0500-D) and 1% Penicillin-Streptomycin. They were transfected with the calcium phosphate method with 150 ng of the plasmids expressing Liprin-α3 (or of RIM1α if Liprin-α was not included), and additional plasmids were added at 1:1 molar ratios (except for Ca_V_2.1, which was added at 0.67:1). 16-20 hours after transfection, cells were either fixed in 4% PFA in PBS and subsequently mounted for imaging or they were used for FRAP. For experiments in Fig. 6j-k, to assess the presence of Munc13-1 (not shown in the figure), cells were blocked and permeabilized in a blocking solution (PBS, 3% BSA, 0.1% Triton X-100) for 1 hour at room temperature, followed by incubations with primary (rabbit anti Munc13-1; A72, 1:500; RRID: AB_887733) and secondary (goat anti-rabbit Alexa Fluor 488; S5; 1:500, RRID: AB_2576217) antibodies overnight at 4 °C in blocking solution. Three 5-minute washes with PBS were performed between steps. All images were acquired in confocal mode using a Leica SP8 Confocal/STED 3X microscope, using an oil-immersion 63x objective (1.44 NA). Images of 2500 µm x 2500 µm or 1850 µm x 1850 µm (pixel size 240 nm x 240 nm) containing multiple transfected cells were taken. Pearson’s correlation was used to assess colocalization of proteins in cells expressing all proteins of interest for any given experiment using the “*Coloc2*” built-in plugin of ImageJ/Fiji. Cells in which the fluorescent signal of at least one of the proteins was covering most of the cell area were not included. In experiments with mE-Ca_V_2.1, cerulean was imaged last to prevent photoconversion of mEOS before other fluorophores were acquired. For FRAP, movies of one single transfected cell were taken at room temperature. Cells were maintained in HEK medium during image acquisition. Single condensates were photobleached using a 405 nm wavelength laser followed by image acquisition at 1 Hz in confocal mode at room temperature, as we described before^20^. Regions of interest were drawn over pre-bleached structures and the percentage of intensity recovered was plotted as a function of time. Each individual trace was normalized to its maximum and minimum intensity values and plotted in Fig. 6f. The maximum value after photobleaching of the resulting normalized trace within the first 120 seconds of recovery was used to calculate the percentage of recovery (Fig. 6g). Only for representative images, a smooth filter was added, brightness and contrast were linearly adjusted, and images were interpolated. These adjustments were done either equally across conditions (Figs. 3, 6b and 6k), or unequally to reach a similar level of signal intensity between condensates before bleaching (Fig. 6e). Quantifications were done on original images without adjustments. pCMV RIM1α-tdTomato (rat, p1010) and pCMV RIM1α-cerulean (rat, p1011) follow the same cDNA sequence variants and numbering as described for p592 above. The fluorescent tags were placed between E_1379_ and S_1380_ and flanked by exogenous linkers (p1010:…E_1379_ – AAAAA-V_2_….K_476_-AAGGAP- S_1380_… with V_2_ to K_476_ referring to tdTomato residues according to GenBank: LC311026.1; p1011:…E_1379_ -AAA-V_2_…K_239_-GAP-S_1380_… with V_2_ to K_239_ referring to Cerulean residues according to GenBank: ACO48272.1). pCMV Cerulean- Liprin-α3 (rat, p471) and pCMV mVenus-Liprin-α3 (rat, p472) are the same sequence variant and numbering as outlined for p576 above and have been described before^20^. The Cerulean or mVenus tags were placed at the N-terminus flanked by linker sequences (sequence: M-AAA- V_2_…K_239_-GAPS-C_3_…C_1192_, V_2_…K_239_ refer to Cerulean for p471 or mVenus for p472 according to GenBank: ACO48272.1 and GenBank: AAZ65844.1, respectively). pCMV HA-Liprin-α3 (mouse, p470) corresponds to GenBank: NP_084017.2 with a T714A point mutation and the HA tag was positioned at the N-terminus (sequence: M-YPYDVPDYA-M_1_…C_1194_). pCMV Cerulean-RIM-BP2 (rat, p1009) and pCMV RIM-BP2 (rat, p205) correspond in sequence and numbering to XP_038945062.1. The cerulean tag was placed at the N-terminus between M_1_ and R_2_ and flanked by short linkers (sequence: M_1_-AAA- V_2_…K_239_-GAP-R_2_…P_1068_; V_2_…K_239_ refer to Cerulean according to GenBank: ACO48272.1). pCMV mEOS-Ca_V_2.1 (mouse, p772) was produced based on a previous expression vector^50^ and numbering and sequence correspond to GenBank: AAW56205.1. It has an mEos3.2 tag between V_27_ and G_28._ that is flanked by short linkers (sequence: M_1_… V_27_ -AS-S_2_…R_226_- ACR- G_28_…C_2369_). S_2_…R_226_ refer to mEos3.2 as described^93^ (https://www.fpbase.org/protein/meos32/). mEos3.2 was obtained as a gift from Michael Davidson and Tao Xu (Addgene plasmid # 54550; http://n2t.net/addgene:54550; RRID:Addgene_54550). pcDNA Munc13-1 (rat, p202) corresponds in sequence and numbering to NCBI Reference Sequence NP_074052.2 without G_1415_TLLRKHGKGLEKGRVKLPSHSD_1437_ and V_1533_HGGKGTRFTLSEDVCPEM_1551_; see NCBI Reference Sequence: XP_017168369.1 for the same splice variant). pcDNA3.1 Dynamin 1 (human, p880) was a gift from Sandra Schmid and was obtained from Addgene (Addgene plasmid #34682; http://n2t.net/addgene:34682; RRID:Addgene_34682). It contains a partial N-terminal HA-tag and a sequence that corresponds to PDB: 7AX3_A (sequence: ME-YDVPDYA-H-M_1_…L_864_). pCMV HA-PTPσ (human, p844, obtained from Jaewon Ko with a modified pDisplay backbone) and pCMV tdTomato-HA-PTPσ (human, p889) follow the same sequence variant and numbering as outlined for p846 above. The N-terminal signal peptide was substituted by a mouse IgΚ leader sequence and followed by the corresponding tag and a short exogenous linker (p844: METDTLLLWVLLLWVPGSTGD*-*YPYDVPDYA-GAQPARS-E_30_…T_1501_; p889: METDTLLLWVLLLWVPGSTGD-YPYDVPDYA-GAQPARS- V_2_…K_476_-RS-E_30_…T_1501_, V_2_ to K_476_ refer to tdTomato residues according GenBank: LC311026.1). pCMV PTPẟ (mouse, p848, in a modified pDisplay vector) was produced from a plasmid containing a PTPδ open reading frame obtained from Jaewon Ko (pcDNA myc-PTPδ^89^). The numbering and the sequence of p848 correspond to GenBank: XP_036019750.1 but with two variants (without amino acids V_51_ and E_189_-I_191_, see NCBI Reference Sequence XP_036019737.1). The signal peptide was replaced with a mouse IgΚ leader sequence followed by a short exogenous linker (sequence: (METDTLLLWVLLLWVPGSTGD-GAPGPV- E_28_-T_1921_). pCMV LAR (human, p849) was cloned from a plasmid obtained from the Harvard Plasmid Repository expressing human LAR identical to the sequence in GenBank: AAH48768.1.

### Correlative light-electron microscopy

HEK293T cells were plated on 25 mm, #2 photo-etched, gridded coverslip in HEK medium, transfected as described for protein condensate experiments above, and fixed in 2.5% glutaraldehyde in PBS buffer (pH 7.4) for 30 minutes at room temperature 16-20 hours after transfection. Fluorescent and wide-field images of transfected cells expressing the proteins of interest were acquired using either a Nikon Ti inverted microscope equipped with Yokagawa CSU-W1 spinning disc confocal system and a 40x objective (0.75 NA) or a Leica SP8 Confocal/STED 3X microscope, using an oil-immersion 63x objective (1.44 NA). After acquiring images, samples were washed 3 times for 5 minutes each with ice cold 100 mM PIPES, pH 7.4 and then incubated in Staining Solution I (SSI; consisting of 1% OsO4, 1.25% potassium hexacyanoferrate in 100 mM PIPES, pH 7.4,) for 2 hours followed by Staining Solution II (prepared by diluting SSI 100 times in a solution of 1% tannic acid) for 30 minutes, and, finally, in 1% uranyl acetate overnight. The samples were kept at 4 °C and protected from light during all the staining steps. Three 5-minute washes with ice-cold mili-Q water were performed between steps. Samples were dehydrated with increasing concentrations of ethanol (30%, 50%, 70%, 90%, 100%, ice-cold), followed by two washes in 100% ice-cold acetone, embedded in epoxy resin, and baked at 60 °C for 36 hours. The coverslip was removed from the resin by sequential immersions into liquid nitrogen and boiling water. 50 nm thick sections from blocks containing grid areas with the cells of interest were cut with a Leica EM UC7 ultramicrotome. A JEOL 1200EX transmission electron microscope equipped with an AMT 2k CCD camera was used for image acquisition of the target cells (identified by their position on the gridded coverslips and by their shape matching that of the wide field image). Fluorescent and electron microscopy images were aligned using the BigWarp plugin (ImageJ/Fiji). The electron micrograph was used as fixed image and arbitrary references (such as the nucleus or the plasma membrane) were used for alignment. The alignment for each cell was conducted multiple times, using each time different features as landmarks, to independently confirm the alignement. For representative images, a smooth filter was added to fluorescent images, and brightness and contrast in all images were linearly adjusted and interpolated.

### Electrophysiology of cultured neurons

Whole-cell patch clamp recordings of cultured hippocampal neurons plated on 12 mm, #1.0 or #1.5 glass coverslips were done as described before at DIV15 to 16^20,21,26,34,50,62^. Glass pipettes (resistance of 1.5 – 4 MΩ) were filled with an intracellular solution containing (in mM) 120 Cs-methanesulfonate, 10 EGTA, 2 MgCl_2_, 10 HEPES-CsOH (pH 7.4), 4 Na_2_-ATP, and 1 Na-GTP for excitatory transmission; and 40 CsCl, 90 K-Gluconate, 1.8 NaCl, 1.7 MgCl_2_, 3.5 KCl, 0.05 EGTA, 10 HEPES, 2 MgATP, 0.4 Na_2_-GTP, 10 phosphocreatine, CsOH (pH 7.4) for inhibitory transmission. Neurons were clamped at -70 mV (or at +30 mV for NMDAR-EPSCs) and series resistance was compensated to ∼5 MΩ. Any recording with series resistance >15 MΩ before compensation at any given point during acquisition was discarded. Recordings were done at room temperature in extracellular solution containing (in mM) 140 NaCl, 5 KCl, 1.5 CaCl_2_, 2 MgCl_2_, 10 HEPES (pH 7.4) and 10 Glucose. For mEPSCs, mIPSCs and sucrose-evoked release, the extracellular solution was supplemented with 1 µM TTX, 50 µM D-AP5 and either 50 µM picrotoxin (for EPSCs) or 20 µM CNQX (for IPSCs). For electrically evoked currents, the extracellular solution was supplemented with 20 µM CNQX and either 50 µM D-AP5 (for IPSCs) or 20 mM PTX (for NMDAR-EPSCs). Electrical stimulation was applied using a bipolar electrode custom-made from Nichrome wire. 500 mM hypertonic sucrose was locally applied with a pump for 10 seconds at a flow rate of 10 µl per minute, and the integral of the first 10 seconds of the response was used to estimate the RRP. A Multiclamp 700B amplifier and a Digidata 1550 digitizer were used, sampling at 10 kHz and filtering at 2 kHz. Data were analyzed using pClamp. Data were acquired and analyzed by an experimenter blind to genotype.

### Electron microscopy of cultured neurons

Electron microscopy was performed as before^20,21,26,34,50,62,63^. Briefly, DIV15 neurons grown on 0.12 mm thick, 6 mm diameter Matrigel-coated sapphire coverslips were transferred to an extracellular solution containing (in mM) 140 NaCl, 5 KCl, 1.5 CaCl_2_, 2 MgCl_2_, 10 glucose, CNQX (20 mM), D-AP5 (50 mM), PTX (50 mM), 10 Hepes (pH 7.4, ∼310 mOsm) and high-pressure frozen with a Leica EM ICE freezer. The samples were then freeze substituted using an acetone solution supplemented with 1% osmium tetroxide, 1% glutaraldehyde, and 1% H_2_O following this protocol: -90 °C for 24 hours, 5 °C per hour to -20 °C, -20 °C for 12 hours, and 10 °C per hour to 20 °C. Next, samples were embedded in epoxy resin and baked for 48 hours at 60 °C. Next, the sapphire coverslip was removed from the resin block by sequential immersions into liquid nitrogen and boiling water. The process was repeated until the coverslip detached.

Then, the resin block containing the neurons was divided into four pieces and each piece was mounted on a stub. A Leica EM UC7 ultramicrotome was used for sectioning at 50 nm, and sections were collected on a nickel slot grid (2 x 1 mm) with a carbon coated formvar support film. The sections were counterstained with 2% lead acetate solution for 10 seconds, followed by rinsing with distilled water. Images were taken with a JEOL 1200EX transmission electron microscope equipped with an AMT 2k CCD camera. Synapses containing a cloud of vesicles and with an evident, well-preserved synaptic cleft were selected for quantification. A MATLAB macro, provided by M. Verhage and J Broeke was used for analyses. Docked vesicles were defined as those in contact with the presynaptic plasma membrane apposed to the PSD and with no white space between the electron-dense vesicular and target membranes. Bouton size is expressed as the area within the perimeter of a bouton in a cross section. Brightness and contrast were adjusted for representative examples to match appearance. Data were acquired and analyzed by an experimenter blind to condition.

### Statistics

Data are shown as mean ± SEM unless noted otherwise. Violin plots are shown when the number of data points for at least one of the groups was >100. Significance was assessed using parametric (t-test or one-way ANOVA) or non-parametric (Mann-Whitney U or Kruskal-Wallis) tests depending on whether assumptions of normality and homogeneity of variances were met (assessed using Shapiro or Levene’s tests, respectively). Tukey-Kramer or Holm corrections for multiple testing were applied. Post-hoc comparisons between all groups were performed, and only significance relative to the respective conditional knockout group (or to the control group in Fig. 3c, the “no PTP” group in Fig. 6c+d, and LAR in Fig. 6g) is reported in figures. For paired pulse ratios, a two-way ANOVA was used followed by Dunnett post hoc test. A Chi-square test was used to assess mouse survival ratios. Data were analyzed by an experimenter blind to the condition/genotype.

---

**Supplemental figure 1.**
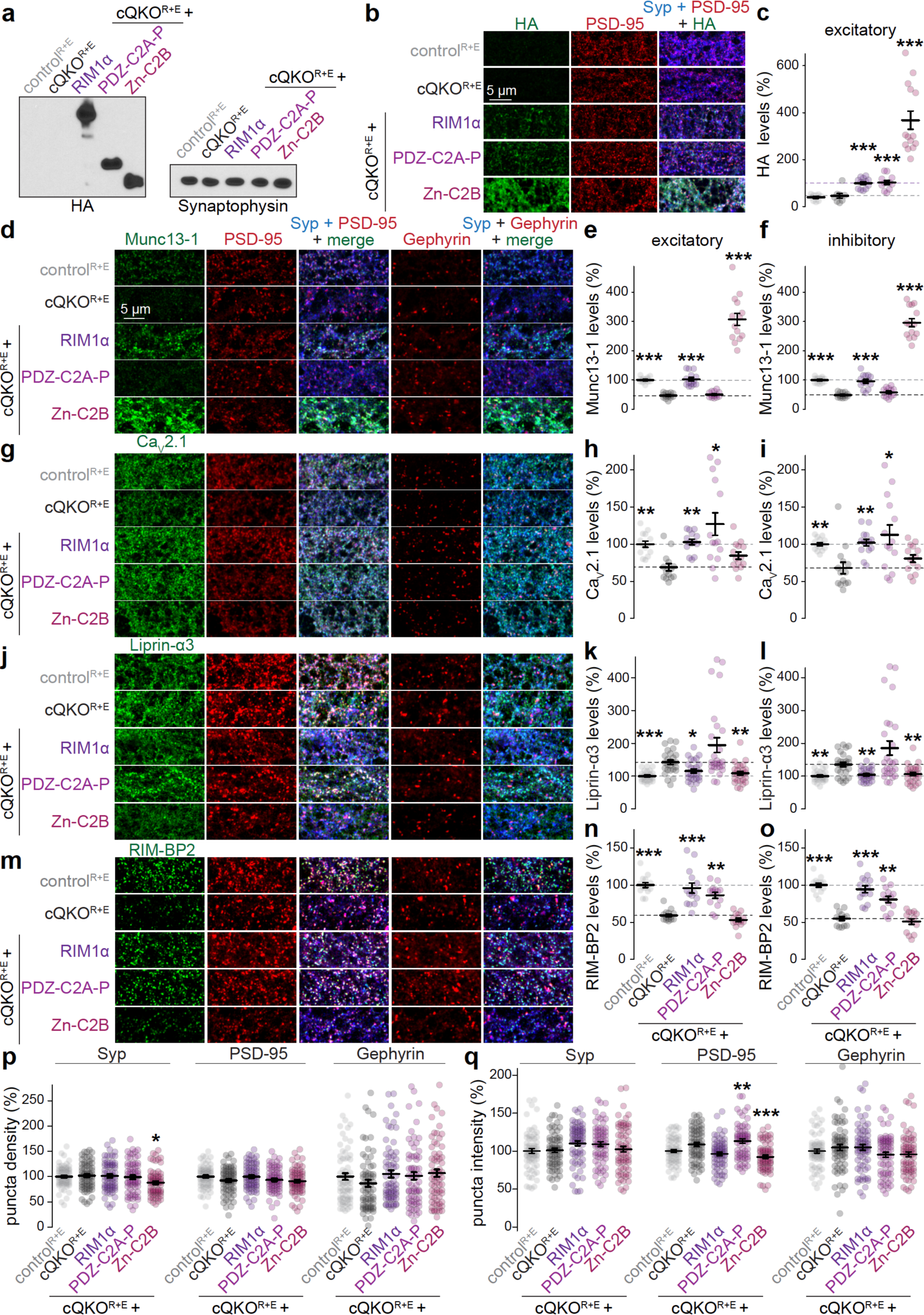
Assessment of RIM1 constructs with Western blot and confocal microscopy (**a**) Western blot to assess expression of RIM1α, PDZ-C2A-P and Zn-C2B in cultured neurons. (**b, c**) Example confocal images (b) and quantification (c) of the average intensity of HA (to detect rescue proteins) at excitatory synapses identified as PSD-95 regions of interest (ROIs). Neurons were stained for HA, PSD-95 and Synaptophysin (Syp). Intensity is normalized to the average cQKO^R+E^ + RIM1α per culture. Levels at inhibitory synapses were not assessed due to incompatibility of HA and Gephyrin antibodies; control^R+E^ 9 images/3 independent cultures, cQKO^R+E^ 8/3, cQKO^R+E^ + RIM1α 13/3, cQKO^R+E^ + PDZ-C2A-P 12/3, cQKO^R+E^ + Zn-C2B 14/3. (**d-o**) Example confocal images and quantification of the average fluorescence intensity levels at excitatory and inhibitory synapses of Munc13-1 (d-f), Ca_V_2.1 (g-i), Liprin-α3 (j-l) and RIM-BP2 (m-o). Neurons were stained for a protein of interest (Munc13-1, Ca_V_2.1, Liprin-α3 or RIM-BP2), postsynaptic markers (PSD-95 and Gephyrin), and Synaptophysin. Excitatory synapses were defined as PSD-95 ROIs and inhibitory synapses as Gephyrin ROIs. Data are normalized to the average control^R+E^ per culture, dotted lines mark the levels of cQKO^R+E^ (black) or control^R+E^ (gray); d-f, control 14/3, cQKO^R+E^ 14/3, cQKO^R+E^ + RIM1α 14/3, cQKO^R+E^ + PDZ-C2A-P 13/3, cQKO^R+E^ + Zn-C2B 14/3; g-i, 14/3 each; j-l, 26/6 each; m-o, 14/3 each. (**p, q**) Quantification of Synaptophysin, PSD-95 and Gephyrin puncta densities (p) and of their fluorescence intensities (q) normalized to the average control^R+E^ per culture. Small changes in Synaptophysin and PSD-95 in some conditions do not confound the conclusion that independent assembly pathways recruit Munc13-1 and Ca_V_2.1; control^R+E^ 68/6, cQKO^R+E^ 68/6, cQKO^R+E^ + RIM1α 68/6, cQKO^R+E^ + PDZ-C2A-P 67/6, cQKO^R+E^ + Zn-C2B 68/6. Data are mean ± SEM; *p < 0.05, **p < 0.01, ***p < 0.001 compared to cQKO^R+E^ as determined by Kruskal-Wallis followed by Holm multiple comparisons post hoc tests in c, e, f, h, i, k, l, n, o, p (Synaptophysin and Gephyrin) and q, or by a one-way ANOVA followed by Tukey-Kramer multiple comparisons post hoc tests in p (PSD-95).

**Supplemental figure 2.**
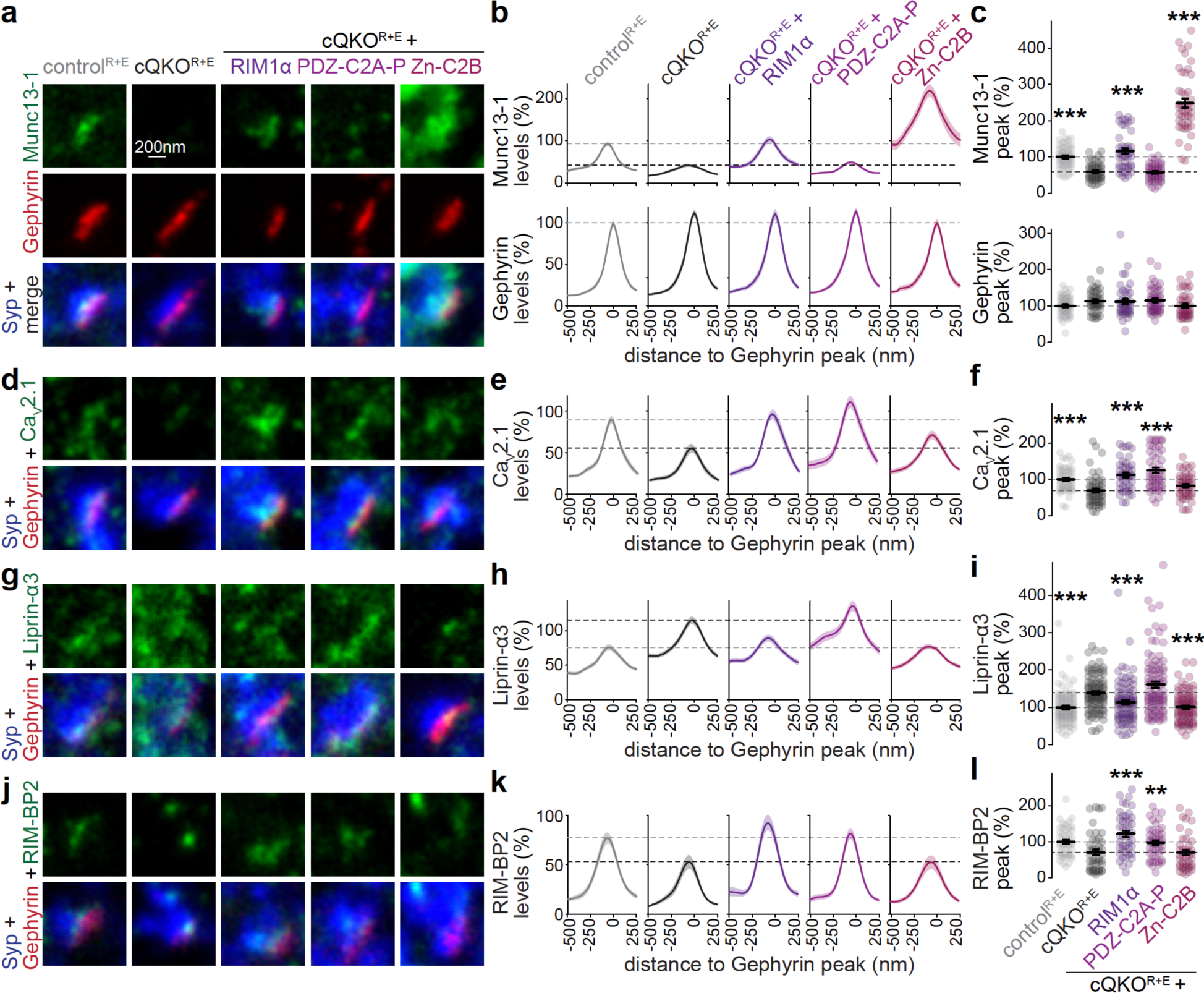
Assessment of inhibitory synapses by STED microscopy after RIM rescue (**a-l**) Example STED images, average line profiles and quantification of the peak intensity of Munc13-1 and Gephyrin (a-c), Ca_V_2.1 (d-f), Liprin-α3 (g-i) and RIM-BP2 (j-l) at inhibitory side- view synapses identified by Synaptophysin (Syp) and Gephyrin. Analyses were performed on the experiment shown in Fig. 2f-q as the neurons were co-stained for Gephyrin. A line profile (750 nm x 250 nm) was positioned perpendicular to the center of the elongated Gephyrin object and profiles of all synapses were aligned to the Gephyrin peak. The maximum value of each individual profile was used to calculate the peak. Dotted lines mark the levels of cQKO^R+E^ (black) or control^R+E^ (gray), line profiles and peak intensities are normalized to the average control^R+E^ per culture; a-c, control^R+E^ 45 synapses/3 independent cultures, cQKO^R+E^ 52/3, cQKO^R+E^ + RIM1α 44/3, cQKO^R+E^ + PDZ-C2A-P 49/3, cQKO^R+E^ + Zn-C2B 45/3; d-f, control^R+E^ 49/3, cQKO^R+E^ 52/3, cQKO^R+E^ + RIM1α 44/3, cQKO^R+E^ + PDZ-C2A-P 51/3, cQKO^R+E^ + Zn-C2B 47/3; g-i, control^R+E^ 94/6, cQKO^R+E^ 97/6, cQKO^R+E^ + RIM1α 94/6, cQKO^R+E^ + PDZ-C2A-P 95/6, cQKO^R+E^ + Zn-C2B 91/6; j-l, control^R+E^ 46/3, cQKO^R+E^ 46/3, cQKO^R+E^ + RIM1α 45/3, cQKO^R+E^ + PDZ-C2A-P 47/3, cQKO^R+E^ + Zn-C2B 44/3. Data are mean ± SEM; **p < 0.01, ***p < 0.001 as determined by Kruskal-Wallis followed by Holm multiple comparisons post hoc tests.

**Supplemental figure 3.**
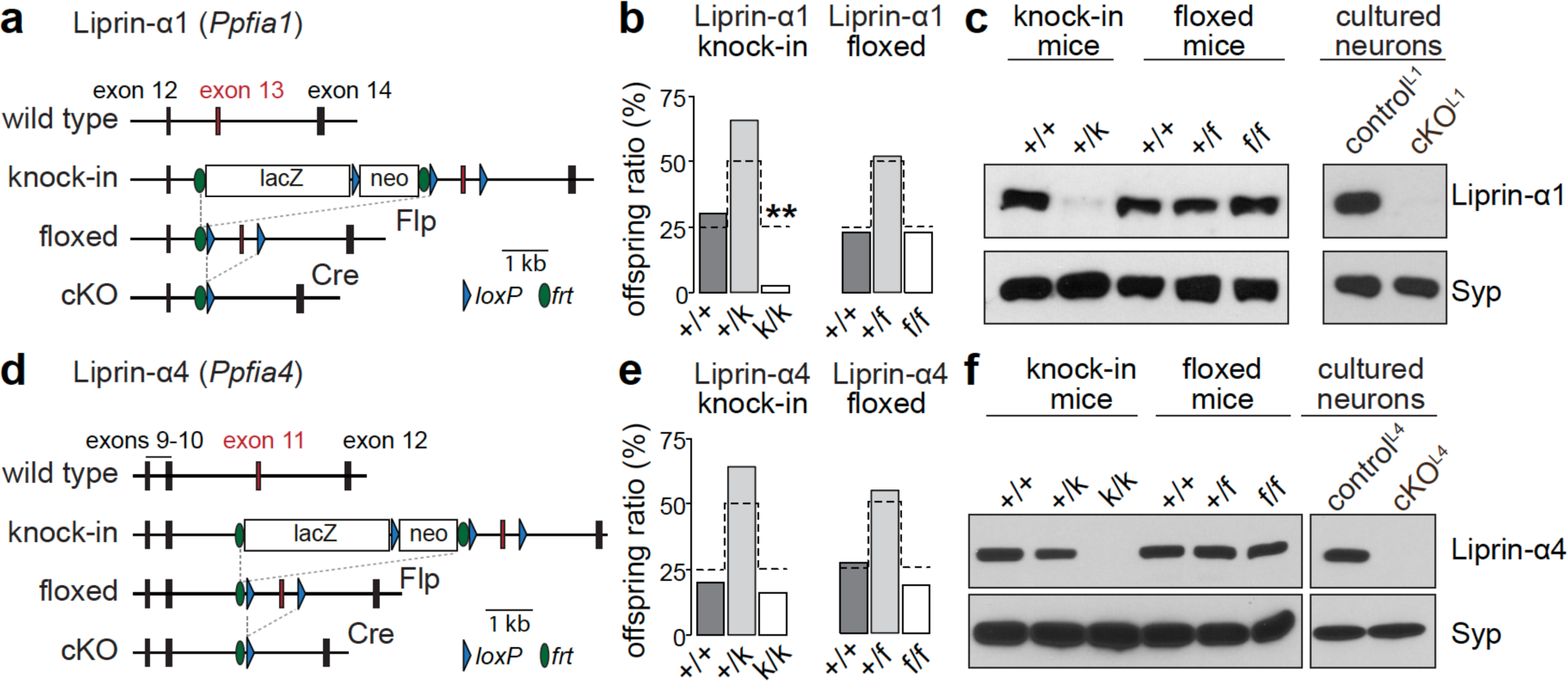
Liprin-α1 and -α4 mutant alleles (a) Diagram outlining the gene targeting strategy of *Ppfia1* to remove Liprin-α1. The knock-in allele containing loxP sites flanking exon 13 (numbering follows Ensembl ENSMUST00000182226.8) was generated by homologous recombination, chimeric founders were used to establish the knock-in line and subsequently crossed to Flp-transgenic mice^88^ to generate the floxed allele. (b) Offspring ratios from Liprin-α1 knock-in (k) or floxed (f) heterozygous breeding pairs. Dotted lines show expected Mendelian ratios; Liprin-α1 knock-in 26 mice/5 litters; Liprin-α1 floxed 35/5. (c) Western blots of whole brain homogenates of wild type, heterozygous and homozygous littermate Liprin-α1 mice before (knock-in) or after Flp recombination (floxed), or of lysates from hippocampal cultures of Liprin-α1 floxed mice infected with a lentivirus expressing Cre (cKO^L1^) or a recombination deficient truncation of Cre (control^L1^). (**d-f**) Same as a-c but for *Ppfia4* to remove Liprin-α4, exon 11 is flanked by loxP sites (numbering follows Ensembl ENSMUST00000168515.8); e, Liprin-α4 knock-in 25 mice/4 litters; Liprin-α floxed 44/5. Data in b and e are shown as observed offspring ratios. **p < 0.01 as determined by Chi-square tests comparing obtained ratios with expected Mendelian ratios.

**Supplemental figure 4.**
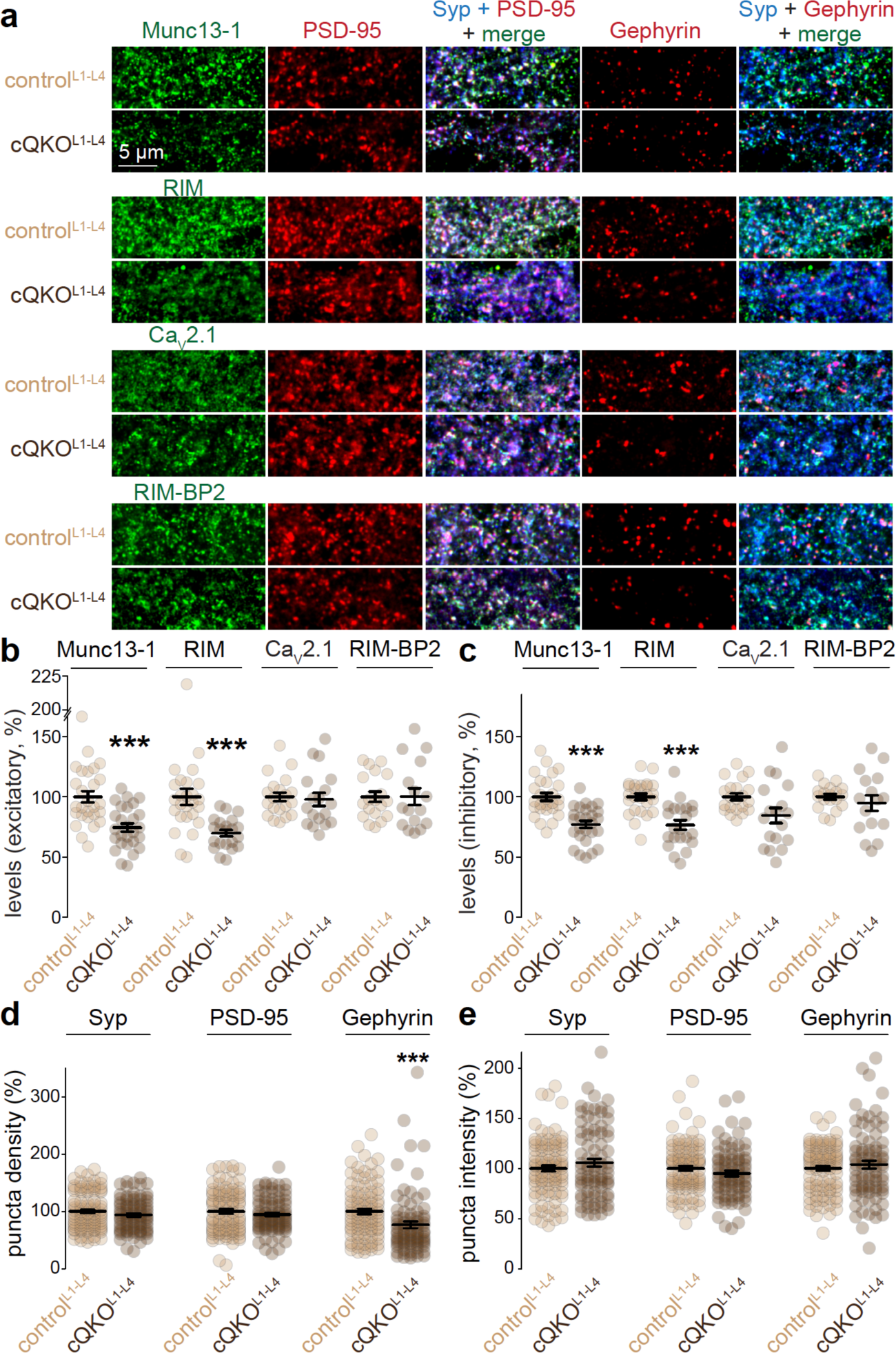
Confocal microscopic assessment of synaptic protein levels after ablation of Liprin-α1 to -α4 (**a-c**) Example confocal images (a) and quantification of fluorescence intensity levels at excitatory and inhibitory synapses (b, c). Neurons were stained for a protein of interest (Munc13-1, RIM, Ca_V_2.1 or RIM-BP2), two postsynaptic markers (PSD-95, excitatory, and Gephyrin, inhibitory), and Synaptophysin (Syp). Data are normalized to the average control^L1-L4^ per culture; Munc13-1, control^L1-L4^ 27 images/3 independent cultures, cQKO^L1-L4^ 27/3; RIM, control^L1-L4^ 26/3, cQKO^L1-L4^ 22/3; Ca_V_2.1, control^L1-L4^ 21/3, cQKO^L1-L4^ 18/3; RIM-BP2, control^L1-L4^19/3, cQKO^L1-L4^ 16/3. (**d, e**) Quantification of Synaptophysin, PSD-95 and Gephyrin puncta densities (d) and of their fluorescence intensities (e) normalized to the average control^L1-L4^ per culture. The ∼20 % reduction in inhibitory synapses is unlikely to fully account for the ∼60% decrease in mIPSC frequency and the ∼45% decrease in sucrose-evoked responses (Fig. 4w+x and Supplemental fig. 5q+r); control^L1-L4^ 93/3, cQKO^L1-L4^ 83/3. Data are mean ± SEM; ***p < 0.001 compared to cQKO^L1-L4^ as determined by Mann-Whitney U tests in b (RIM and RIM-BP2), c (RIM, RIM-BP2, and Ca_V_2.1), d, and e, or by Student’s t-tests in b (Munc13-1 and Ca_V_2.1) and c (Munc13-1).

**Supplemental figure 5.**
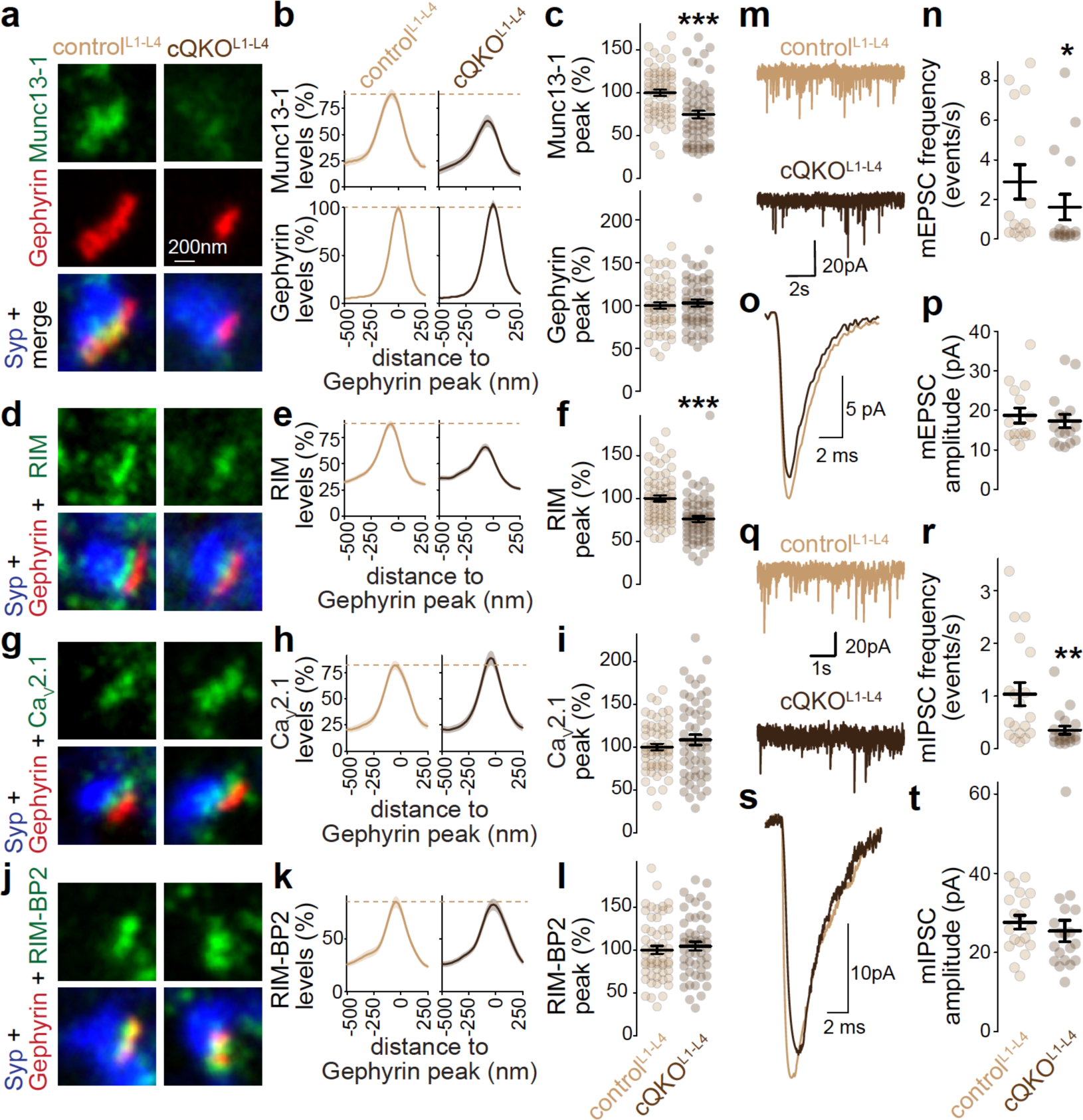
Assessment of inhibitory synapses by STED microscopy and of spontaneous transmission after Liprin-α ablation (**a-l**) Example STED images, average line profiles and quantification of the peak intensity of Munc13-1 and Gephyrin (a-c), RIM (d-f), Ca_V_2.1 (g-i) and RIM-BP2 (j-l) at inhibitory side-view synapses identified by Synaptophysin (Syp) and Gephyrin. Analyses were performed on the experiment shown in Fig. 4c-n as the neurons were co-stained for Gephyrin. Dotted lines in line profile plots mark the levels of control^L1-L4^, line profiles and peak intensities are normalized to the average control^L1-L4^ per culture; a-c, control^L1-L4^ 60 synapses/3 independent cultures, cQKO^L1-L4^ 61/3; d-f, control^L1-L4^ 68/3, cQKO^L1-L4^ 65/3; g-i, control^L1-L4^ 58/3, cQKO^L1-L4^ 56/3; j-l, control^L1-L4^57/3, cQKO^L1-L4^ 51/3. (**m, n**) Example traces (m) of spontaneous miniature excitatory postsynaptic current (mEPSC) recordings and quantification (n) of mEPSC frequency; control^L1-L4^ 15 cells/3 independent cultures, cQKO^L1-L4^ 16/3. (**o, p**) Example traces (o) of an averaged mEPSC from a single cell and quantification of the mEPSC amplitude (p), N as in m+n. (**q-t**) As in m-p but for mIPSCs; 19/3 each. Data are mean ± SEM; ***p < 0.001 compared to cQKO^L1-L4^ as determined by Mann-Whitney U tests (c, f, i, n, p, r and t) or by a Student’s t-tests (l).

**Supplemental figure 6.**
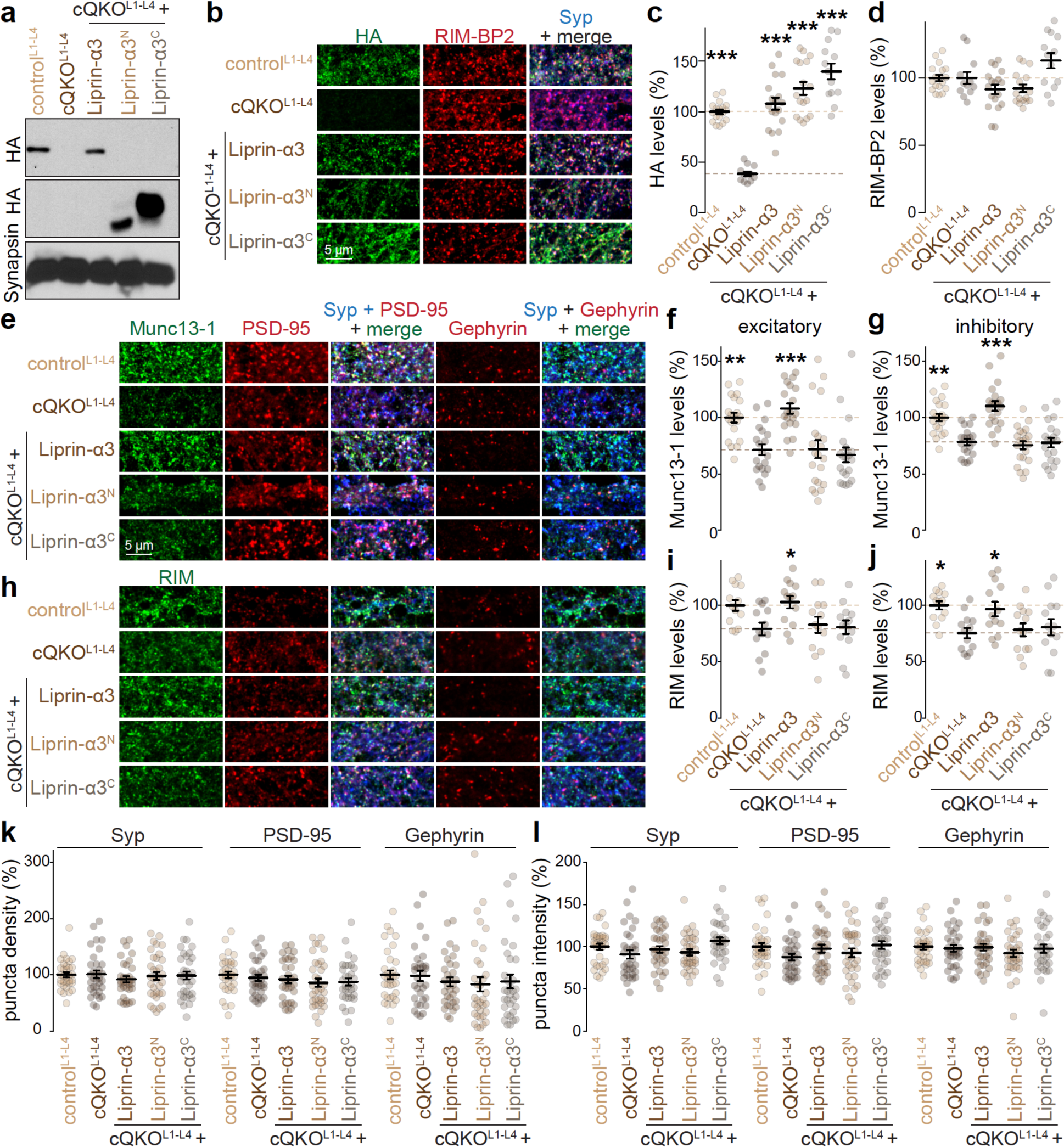
Assessment of Liprin-α rescue constructs with Western blot and confocal microscopy (a) Western blot to assess expression of Liprin-α3, Liprin-α3^N^ and Liprin-α3^C^; note that control^L1-L4^ has an HA signal because of the lentiviral expression of HA-tagged Liprin-α3. (**b-d**) Example confocal images (a) and quantification of fluorescence intensity levels (b+c) of HA to detect Liprin-α3 and of RIM-BP2 at synapses defined as Synaptophysin (Syp) ROIs. Intensity is normalized to the average control^L1-L4^ per culture, dotted lines mark the levels of cQKO^L1-L4^ (dark brown) or control^L1-L4^ (light brown); control^L1-L4^ 18 images/5 independent cultures, cQKO^L1-L4^ 12/5, cQKO^L1-L4^ + Liprin-α3 18/5, cQKO ^L1-L4^ + Liprin-α3^N^ 17/5, cQKO^L1-L4^ + Liprin-α3^C^ 13/5. (**e-j**) Example confocal images and quantification of fluorescence intensity levels at excitatory and inhibitory synapses of Munc13-1 (e-g) and RIM (h-j). Neurons were stained for a protein of interest (Munc13-1 or RIM), postsynaptic markers (PSD-95 and Gephyrin), and Synaptophysin. Data are normalized to the average control^L1-L4^ per culture, dotted lines mark the levels of cQKO^L1-L4^ (dark brown) or control^L1-L4^ (light brown); e-g, control^L1-L4^ 20/4, cQKO^L1-L4^ 20/4, cQKO^L1-L4^ + Liprin-α3 20/4, cQKO ^L1-L4^ + Liprin-α3^N^ 21/4, cQKO^L1-L4^ + Liprin-α3^C^ 20/4; h-j, 13/3 each. (**k, l**) Quantification of Synaptophysin, PSD-95 and Gephyrin puncta densities (k) and of their fluorescence intensities (l) normalized to the average control^L1-L4^ per culture; control^L1-L4^ 33/4, cQKO^L1-L4^ 33/4, cQKO^L1-L4^ + Liprin-α3 33/4, cQKO ^L1-L4^ + Liprin-α3^N^ 34/4, cQKO^L1-L4^ + Liprin-α3^C^ 33/4. Data are mean ± SEM; *p < 0.05 **p < 0.01, ***p < 0.00 compared to cQKO^L1-L4^ as determined by Kruskal-Wallis followed by Holm multiple comparisons post hoc tests in f and k (Gephyrin and Synaptophysin) and l (Synaptophysin), or by one-way ANOVA followed by multiple comparisons Tukey-Kramer post hoc tests in g, i, j, k (PSD-95) and l (PSD-95 and Gephyrin).

**Supplemental figure 7.**
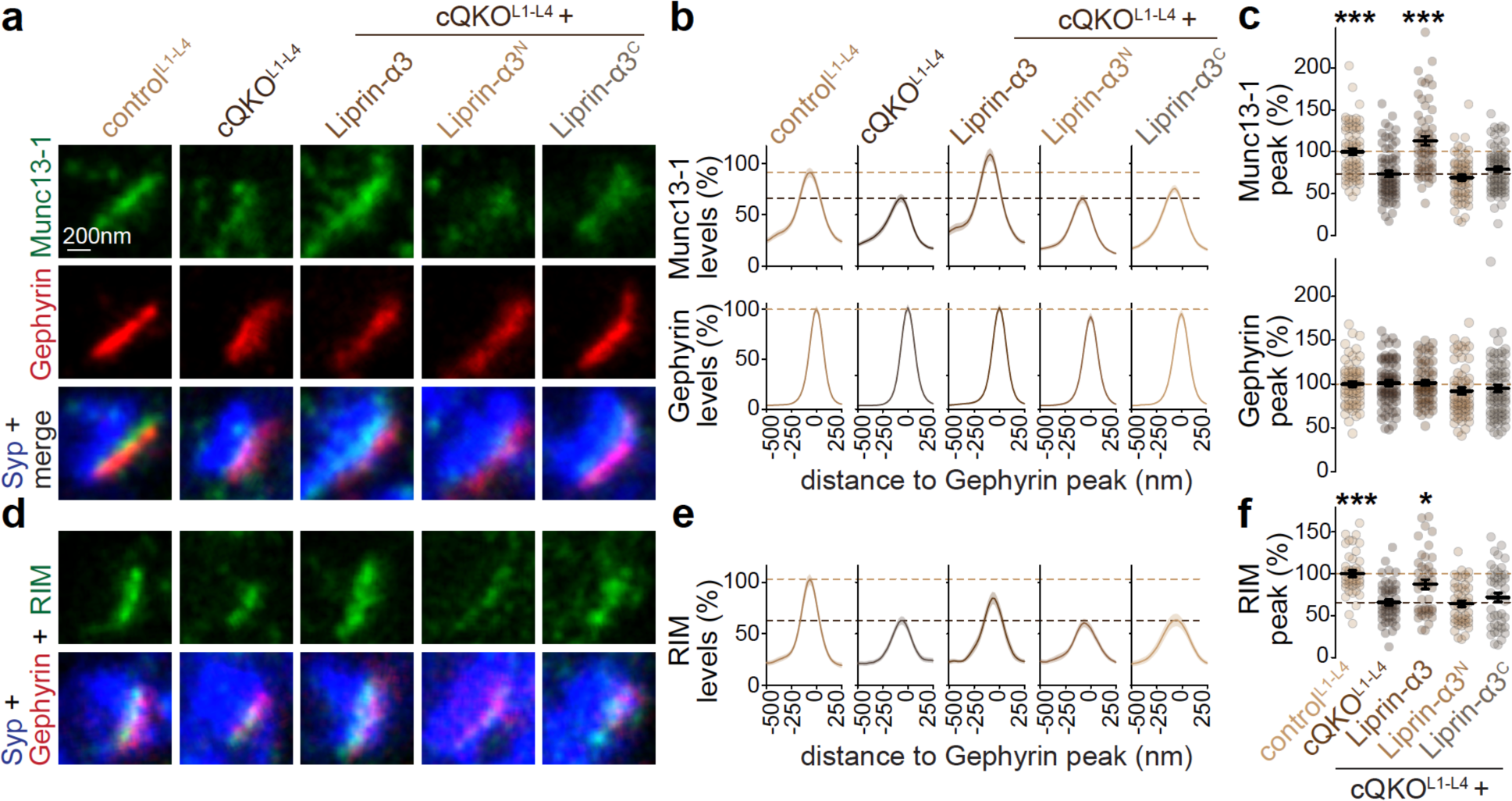
Assessment of inhibitory synapses by STED microscopy after Liprin-α3 rescue (**a-f**) Example STED images, average line profiles and quantification of the peak intensity of Munc13-1 and Gephyrin (a-c), and RIM (d-f) at inhibitory side-view synapses identified by Synaptophysin (Syp) and Gephyrin. Analyses were performed on the experiment shown in Fig. 5b-i as the neurons were co-stained for Gephyrin. Dotted lines mark the levels of cQKO^L1-L4^ (dark brown) or control^L1-L4^ (light brown), line profiles and peak intensities are normalized to the average control^L1-L4^ per culture; a-c, control^L1-L4^ 67 synapses/4 independent cultures, cQKO^L1-L4^ 69/4, cQKO^L1-L4^ + Liprin-α3 61/4, cQKO ^L1-L4^ + Liprin-α3^N^ 60/4, cQKO^L1-L4^ + Liprin-α3^C^ 74/4; d-f, control^L1-L4^ 45/3, cQKO^L1-L4^ 48/3, cQKO^L1-L4^ + Liprin-α3 43/3, cQKO ^L1-L4^ + Liprin-α3^N^ 47/3, cQKO^L1-L4^ + Liprin-α3^C^ 44/4. Data are mean ± SEM; *p < 0.05, ***p < 0.001 compared to cQKO^L1-L4^ as determined by Kruskal-Wallis followed by Holm multiple comparisons post hoc tests.

**Supplemental figure 8.**
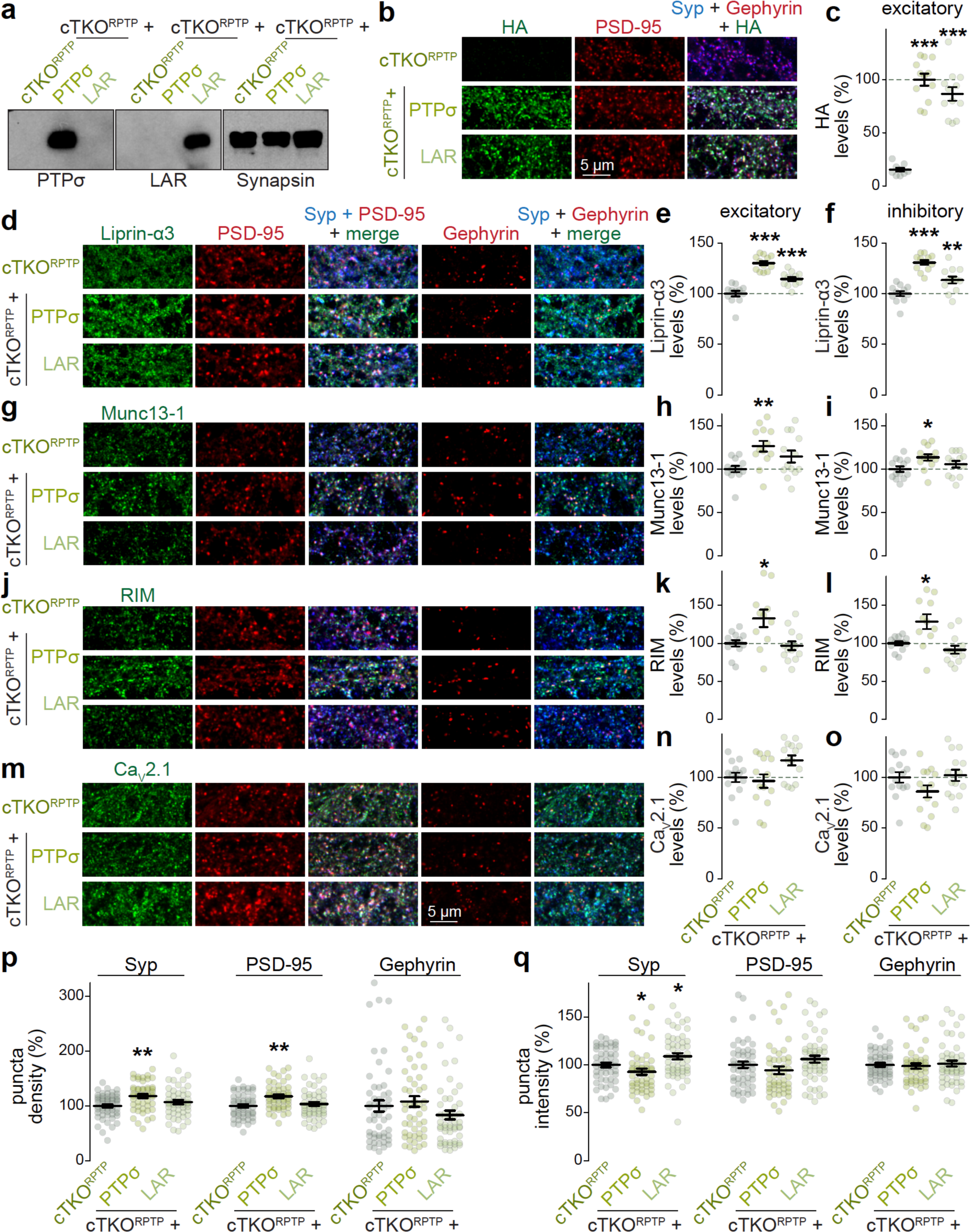
Assessment of LAR-RPTP proteins with Western blot and confocal microscopy (a) Western blot to assess expression of PTPσ and LAR. (**b, c**) Example confocal images and quantification of fluorescence intensity levels of HA to detect PTPσ or LAR at excitatory synapses defined as PSD-95 ROIs. Neurons were stained for HA, PSD-95 and Synaptophysin (Syp). Intensity is normalized to the average cTKO^RPTP^ per culture, dotted lines mark the levels of cTKO^RPTP^ + PTPσ; cTKO^RPTP^ 9 images/3 independent cultures, cTKO^RPTP^ + PTPσ 12/3, cTKO^RPTP^ + LAR 12/3. (**d-o**) Example confocal images and quantification of fluorescence intensity levels at excitatory and inhibitory synapses of Liprin-α3 (d-f), Munc13-1 (g-i), RIM (j-l) and Ca_V_2.1 (m-o). Neurons were stained for a protein of interest (Liprin-α3, Munc13-1, RIM or Ca_V_2.1), postsynaptic markers (PSD-95 and Gephyrin), and Synaptophysin. Data are normalized to the average cTKO^RPTP^ per culture, dotted lines mark the levels of cTKO^RPTP^; d-f, 12/3 each; g-i, 13/3 each; j-l, cTKO^RPTP^ 13/3, cTKO^RPTP^ + PTPσ 11/3, cTKO^RPTP^ + LAR 13/3; m-o, 14/3 each. **(p, q)** Quantification of Synaptophysin, PSD-95 and Gephyrin puncta densities (p) and of their fluorescence intensities (q) normalized to the average cTKO^RPTP^ per culture. Expression of PTPσ resulted in a mild increase in the number of Synaptophysin and PSD-95 puncta, possibly reflecting a synaptogenic effect, and a mild decrease in the intensity of Synaptophysin; cTKO^RPTP^ 52/3, cTKO^RPTP^ + PTPσ 50/3, cTKO^RPTP^ + LAR 52/3. Data are mean ± SEM; *p < 0.05, **p < 0.01, ***p < 0.001 compared to cTKO^RPTP^ as determined by Kruskal-Wallis followed by Holm multiple comparisons post hoc tests for c, e, k, I, p (Gephyrin), and q, or by a one-way ANOVA followed by Tukey-Kramer multiple comparisons post hoc tests for f, h, i, n, o and p (Synaptophysin and PSD-95).

**Supplemental figure 9.**
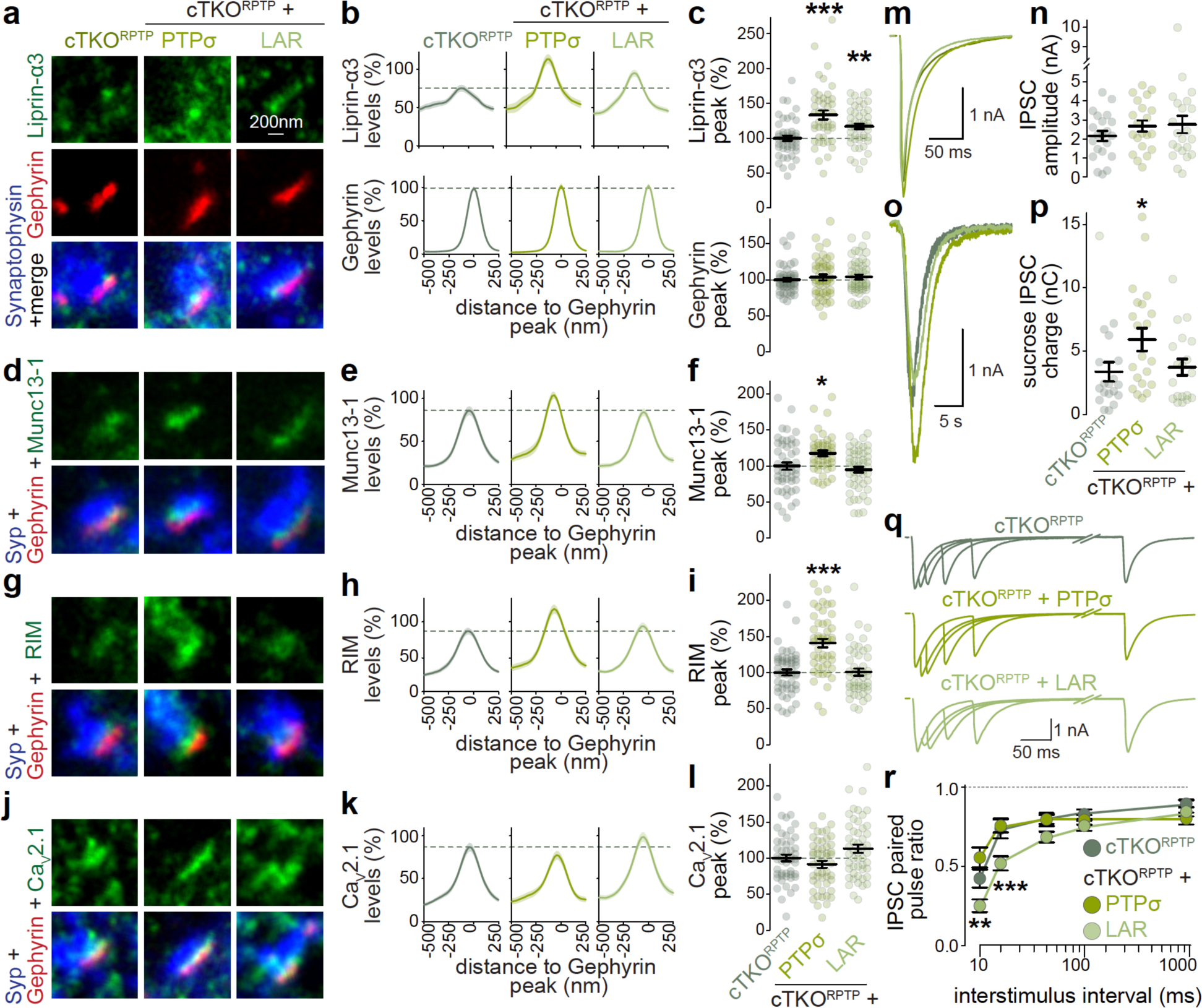
Assessment of inhibitory synapses by STED microscopy and of synaptic transmission after expression of LAR-PTPs in cTKO^RPTP^ neurons (**a-l**) Example STED images, average line profiles and quantification of the peak intensity of Liprin-α3 and Gephyrin (a-c), Munc13-1 (d-f), RIM (g-i) and Ca_V_2.1 (j-l) at inhibitory side-view synapses identified by Synaptophysin (Syp) and Gephyrin. Analyses were performed on the experiment shown in Fig. 7f-q as the neurons were co-stained for Gephyrin. Dotted lines mark the levels of cTKO^RPTP^, line profiles and peak intensities are normalized to the average cTKO^RPTP^ per culture; a-c, cTKO^RPTP^ 48 synapses/3 independent cultures cTKO^RPTP^ + PTPσ 45/3, cTKO^RPTP^ + LAR 50/3; d-f, cTKO^RPTP^ 49/3, cTKO^RPTP^ + PTPσ 45/3, cTKO^RPTP^ + LAR 50/3; g-i, cTKO^RPTP^ 51/3, cTKO^RPTP^ + PTPσ 49/3, cTKO^RPTP^ + LAR 47/3; j-l, cTKO^RPTP^ 49/3, cTKO^RPTP^ + PTPσ 51/3, cTKO^RPTP^ + LAR 50/3. (**m, n**) Example traces (m) and average amplitudes (n) of single action potential-evoked IPSCs. cTKO^RPTP^ 21 cells/3 independent cultures cTKO^RPTP^ + PTPσ 21/3, cTKO^RPTP^ + LAR 22/3. (**o, p**) Example traces (o) and average GABAR-mediated charge transferred in response to hypertonic sucrose superfusion (p); cTKO^RPTP^ 19/3, cTKO^RPTP^ + PTPσ 21/3, cTKO^RPTP^ + LAR 19/3. (**q, r**) Example traces (q) and average IPSC paired pulse ratios (r) at increasing interstimulus intervals to estimate P; N as in m+n. Data are mean ± SEM; *p < 0.05, **p < 0.01, ***p < 0.001 compared to cTKO^RPTP^ as determined by Kruskal-Wallis followed by Holm multiple comparisons post hoc tests (c, f, i, n, and p), by a one-way ANOVA followed by Tukey-Kramer multiple comparisons post hoc tests (l), or by a two- way ANOVA followed by Dunnett multiple comparisons post hoc tests (r).

## References

1. Emperador-Melero, J. & Kaeser, P. S. Assembly of the presynaptic active zone. Curr Opin Neurobiol 63, 95–103 (2020).

2. Südhof, T. C. The Presynaptic Active Zone. Neuron 75, 11–25 (2012).

3. Imig, C. et al. The Morphological and Molecular Nature of Synaptic Vesicle Priming at Presynaptic Active Zones. Neuron 84, 416–431 (2014).

4. Deng, L., Kaeser, P. S., Xu, W. & Südhof, T. C. RIM proteins activate vesicle priming by reversing autoinhibitory homodimerization of Munc13. Neuron 69, 317–331 (2011).

5. Betz, A. et al. Munc13-1 is a presynaptic phorbol ester receptor that enhances neurotransmitter release. Neuron 21, 123–136 (1998).

6. Augustin, I., Rosenmund, C., Südhof, T. C. & Brose, N. Munc13-1 is essential for fusion competence of glutamatergic synaptic vesicles. Nature 400, 457–61 (1999).

7. Richmond, J. E., Weimer, R. M. & Jorgensen, E. M. An open form of syntaxin bypasses the requirement for UNC-13 in vesicle priming. Nature 412, 338–341 (2001).

8. Wu, X. et al. RIM and RIM-BP Form Presynaptic Active-Zone-like Condensates via Phase Separation. Mol Cell 73, 971–984.e5 (2019).

9. Hibino, H. et al. RIM binding proteins (RBPs) couple Rab3-interacting molecules (RIMs) to voltage-gated Ca(2+) channels. Neuron 34, 411–423 (2002).

10. Kaeser, P. S. et al. RIM proteins tether Ca2+ channels to presynaptic active zones via a direct PDZ-domain interaction. Cell 144, 282–95 (2011).

11. Liu, K. S. Y. et al. RIM-binding protein, a central part of the active zone, is essential for neurotransmitter release. Science 334, 1565–9 (2011).

12. Acuna, C., Liu, X. & Südhof, T. C. How to Make an Active Zone: Unexpected Universal Functional Redundancy between RIMs and RIM-BPs. Neuron 91, 792–807 (2016).

13. Dong, W. et al. CAST/ELKS Proteins Control Voltage-Gated Ca2+ Channel Density and Synaptic Release Probability at a Mammalian Central Synapse. Cell Rep 24, 284–293.e6 (2018).

14. Kittel, R. J. et al. Bruchpilot promotes active zone assembly, Ca2+ channel clustering, and vesicle release. Science 312, 1051–4 (2006).

15. Held, R. G., Liu, C. & Kaeser, P. S. ELKS controls the pool of readily releasable vesicles at excitatory synapses through its N-terminal coiled-coil domains. Elife 5, (2016).

16. Liu, C. et al. The Active Zone Protein Family ELKS Supports Ca ^2+^ Influx at Nerve Terminals of Inhibitory Hippocampal Neurons. The Journal of Neuroscience 34, 12289– 12303 (2014).

17. Fenster, S. D. et al. Piccolo, a presynaptic zinc finger protein structurally related to bassoon. Neuron 25, 203–214 (2000).

18. Mukherjee, K. et al. Piccolo and bassoon maintain synaptic vesicle clustering without directly participating in vesicle exocytosis. Proc Natl Acad Sci U S A 107, 6504–6509 (2010).

19. Davydova, D. et al. Bassoon specifically controls presynaptic P/Q-type Ca(2+) channels via RIM-binding protein. Neuron 82, 181–94 (2014).

20. Emperador-Melero, J. et al. PKC-phosphorylation of Liprin-α3 triggers phase separation and controls presynaptic active zone structure. Nat Commun 12, 3057 (2021).

21. Wong, M. Y. et al. Liprin-α3 controls vesicle docking and exocytosis at the active zone of hippocampal synapses. Proc Natl Acad Sci U S A 115, 2234–2239 (2018).

22. Dai, Y. et al. SYD-2 Liprin-alpha organizes presynaptic active zone formation through ELKS. Nat Neurosci 9, 1479–1487 (2006).

23. McDonald, N. A., Fetter, R. D. & Shen, K. Assembly of synaptic active zones requires phase separation of scaffold molecules. Nature 588, 454–458 (2020).

24. Kaufmann, N., DeProto, J., Ranjan, R., Wan, H. & Van Vactor, D. Drosophila liprin-alpha and the receptor phosphatase Dlar control synapse morphogenesis. Neuron 34, 27–38 (2002).

25. Zhen, M. & Jin, Y. The liprin protein SYD-2 regulates the differentiation of presynaptic termini in C. elegans. Nature 401, 371–5 (1999).

26. Wang, S. S. H. et al. Fusion Competent Synaptic Vesicles Persist upon Active Zone Disruption and Loss of Vesicle Docking. Neuron 91, 777–791 (2016).

27. Muller, C. S. et al. Quantitative proteomics of the Cav2 channel nano-environments in the mammalian brain. Proc Natl Acad Sci U S A 107, 14950–14957 (2010).

28. Acuna, C., Liu, X., Gonzalez, A. & Südhof, T. C. RIM-BPs Mediate Tight Coupling of Action Potentials to Ca(2+)-Triggered Neurotransmitter Release. Neuron 87, 1234–1247 (2015).

29. Brockmann, M. M. et al. RIM-BP2 primes synaptic vesicles via recruitment of Munc13-1 at hippocampal mossy fiber synapses. Elife 8, (2019).

30. Lübbert, M. et al. A novel region in the CaV2.1 α1 subunit C-terminus regulates fast synaptic vesicle fusion and vesicle docking at the mammalian presynaptic active zone. Elife 6, (2017).

31. Wong, F. K., Li, Q. & Stanley, E. F. Synaptic vesicle capture by CaV2.2 calcium channels. Front Cell Neurosci 7, (2013).

32. Sheng, Z. H., Westenbroek, R. E. & Catterall, W. A. Physical link and functional coupling of presynaptic calcium channels and the synaptic vesicle docking/fusion machinery. J Bioenerg Biomembr 30, 335–345 (1998).

33. Wu, X. et al. Vesicle Tethering on the Surface of Phase-Separated Active Zone Condensates. Mol Cell 81, 13–24.e7 (2021).

34. Tan, C., Wang, S. S. H., de Nola, G. & Kaeser, P. S. Rebuilding essential active zone functions within a synapse. Neuron 110, 1498–1515.e8 (2022).

35. Vyleta, N. P. & Jonas, P. Loose coupling between Ca2+ channels and release sensors at a plastic hippocampal synapse. Science 343, 665–670 (2014).

36. Rozov, A., Burnashev, N., Sakmann, B. & Neher, E. Transmitter release modulation by intracellular Ca2+ buffers in facilitating and depressing nerve terminals of pyramidal cells in layer 2/3 of the rat neocortex indicates a target cell-specific difference in presynaptic calcium dynamics. J Physiol 531, 807–26 (2001).

37. Rebola, N. et al. Distinct Nanoscale Calcium Channel and Synaptic Vesicle Topographies Contribute to the Diversity of Synaptic Function. Neuron 104, 693–710.e9 (2019).

38. Keller, D. et al. An Exclusion Zone for Ca^2+^ Channels around Docked Vesicles Explains Release Control by Multiple Channels at a CNS Synapse. PLoS Comput Biol 11, (2015).

39. Nakamura, Y. et al. Nanoscale Distribution of Presynaptic Ca2+ Channels and Its Impact on Vesicular Release during Development. Neuron 85, 145–159 (2015).

40. Grauel, M. K. et al. RIM-binding protein 2 regulates release probability by fine-tuning calcium channel localization at murine hippocampal synapses. Proceedings of the National Academy of Sciences 113, 11615–11620 (2016).

41. Böhme, M. A. et al. Active zone scaffolds differentially accumulate Unc13 isoforms to tune Ca2+ channel–vesicle coupling. Nat Neurosci 19, 1311–1320 (2016).

42. Ko, J., Na, M., Kim, S., Lee, J. R. & Kim, E. Interaction of the ERC family of RIM-binding proteins with the liprin-alpha family of multidomain proteins. J Biol Chem 278, 42377– 42385 (2003).

43. Schoch, S. et al. RIM1alpha forms a protein scaffold for regulating neurotransmitter release at the active zone. Nature 415, 321–6 (2002).

44. Wakita, M. et al. Structural insights into selective interaction between type IIa receptor protein tyrosine phosphatases and Liprin-α. Nat Commun 11, (2020).

45. Serra-Pagès, C., Medley, Q. G., Tang, M., Hart, A. & Streuli, M. Liprins, a family of LAR transmembrane protein-tyrosine phosphatase-interacting proteins. J Biol Chem 273, 15611–20 (1998).

46. Xie, X. et al. Structural basis of liprin-α-promoted LAR-RPTP clustering for modulation of phosphatase activity. Nat Commun 11, 169 (2020).

47. Ackley, B. D. et al. The two isoforms of the Caenorhabditis elegans leukocyte-common antigen related receptor tyrosine phosphatase PTP-3 function independently in axon guidance and synapse formation. J Neurosci 25, 7517–7528 (2005).

48. Sakamoto, H. et al. Synaptic weight set by Munc13-1 supramolecular assemblies. Nat Neurosci 21, 41–49 (2018).

49. Reddy-Alla, S. et al. Stable Positioning of Unc13 Restricts Synaptic Vesicle Fusion to Defined Release Sites to Promote Synchronous Neurotransmission. Neuron 95, 1350–1364.e12 (2017).

50. Held, R. G. et al. Synapse and Active Zone Assembly in the Absence of Presynaptic Ca2+ Channels and Ca2+ Entry. Neuron 107, 667–683.e9 (2020).

51. Cao, Y.-Q. et al. Presynaptic Ca2+ channels compete for channel type-preferring slots in altered neurotransmission arising from Ca2+ channelopathy. Neuron 43, 387–400 (2004).

52. Takahashi, T. & Momiyama, A. Different types of calcium channels mediate central synaptic transmission. Nature 366, 156–8 (1993).

53. Dharmasri, P. A., Levy, A. D. & Blanpied, T. A. Differential nanoscale organization of excitatory synapses onto excitatory vs inhibitory neurons. bioRxiv 2023.09.06.556279 (2023) doi:10.1101/2023.09.06.556279.

54. Schnitzbauer, J., Strauss, M. T., Schlichthaerle, T., Schueder, F. & Jungmann, R. Super- resolution microscopy with DNA-PAINT. Nat Protoc 12, 1198–1228 (2017).

55. Jungmann, R. et al. Multiplexed 3D cellular super-resolution imaging with DNA-PAINT and Exchange-PAINT. Nat Methods 11, 313–318 (2014).

56. Ramsey, A. M. et al. Subsynaptic positioning of AMPARs by LRRTM2 controls synaptic strength. Sci Adv 7, (2021).

57. Tang, A.-H. et al. A trans-synaptic nanocolumn aligns neurotransmitter release to receptors. Nature 536, 210–214 (2016).

58. Kershberg, L., Banerjee, A. & Kaeser, P. S. Protein composition of axonal dopamine release sites in the striatum. Elife 11, 2022.08.31.505994 (2022).

59. Dulubova, I. et al. A Munc13/RIM/Rab3 tripartite complex: from priming to plasticity? EMBO J 24, 2839–2850 (2005).

60. Betz, A. et al. Functional interaction of the active zone proteins Munc13-1 and RIM1 in synaptic vesicle priming. Neuron 30, 183–196 (2001).

61. de Jong, A. P. H. et al. RIM C2B Domains Target Presynaptic Active Zone Functions to PIP2-Containing Membranes. Neuron 98, 335–349.e7 (2018).

62. Emperador-Melero, J., de Nola, G. & Kaeser, P. S. Intact synapse structure and function after combined knockout of PTPδ, PTPσ and LAR. Elife 2021.01.17.427005 (2021) doi:10.1101/2021.01.17.427005.

63. Tan, C. et al. Munc13 supports fusogenicity of non-docked vesicles at synapses with disrupted active zones. Elife 11, 2022.04.01.486686 (2022).

64. Liang, M. et al. Oligomerized liprin-α promotes phase separation of ELKS for compartmentalization of presynaptic active zone proteins. Cell Rep 34, 108901 (2021).

65. Zürner, M. & Schoch, S. The mouse and human Liprin-alpha family of scaffolding proteins: genomic organization, expression profiling and regulation by alternative splicing. Genomics 93, 243–253 (2009).

66. Zucker, R. S. & Regehr, W. G. Short-term synaptic plasticity. Annu Rev Physiol 64, 355– 405 (2002).

67. Kaeser, P. S. & Regehr, W. G. The readily releasable pool of synaptic vesicles. Curr Opin Neurobiol 43, 63–70 (2017).

68. Sclip, A. & Südhof, T. C. LAR receptor phospho-tyrosine phosphatases regulate NMDA- receptor responses. Elife 9, (2020).

69. Sclip, A. & Südhof, T. C. Combinatorial expression of neurexins and LAR-type phosphotyrosine phosphatase receptors instructs assembly of a cerebellar circuit. Nat Commun 14, 4976 (2023).

70. Woo, J. et al. Trans-synaptic adhesion between NGL-3 and LAR regulates the formation of excitatory synapses. Nat Neurosci 12, 428–437 (2009).

71. Dunah, A. W. et al. LAR receptor protein tyrosine phosphatases in the development and maintenance of excitatory synapses. Nat Neurosci 8, 458–467 (2005).

72. Prakash, S. et al. Complex interactions amongst N-cadherin, DLAR, and Liprin-alpha regulate Drosophila photoreceptor axon targeting. Dev Biol 336, 10–19 (2009).

73. Takahashi, H. & Craig, A. M. Protein tyrosine phosphatases PTPδ, PTPσ, and LAR: presynaptic hubs for synapse organization. Trends Neurosci 36, 522–534 (2013).

74. Kiyonaka, S. et al. RIM1 confers sustained activity and neurotransmitter vesicle anchoring to presynaptic Ca2+ channels. Nat Neurosci 10, 691–701 (2007).

75. Han, Y., Kaeser, P. S., Südhof, T. C. & Schneggenburger, R. RIM determines Ca^2^+ channel density and vesicle docking at the presynaptic active zone. Neuron 69, 304–16 (2011).

76. Müller, M., Liu, K. S. Y., Sigrist, S. J. & Davis, G. W. RIM controls homeostatic plasticity through modulation of the readily-releasable vesicle pool. J Neurosci 32, 16574–16585 (2012).

77. Andrews-Zwilling, Y. S., Kawabe, H., Reim, K., Varoqueaux, F. & Brose, N. Binding to Rab3A-interacting molecule RIM regulates the presynaptic recruitment of Munc13-1 and ubMunc13-2. J Biol Chem 281, 19720–19731 (2006).

78. Camacho, M. et al. Heterodimerization of Munc13 C2A domain with RIM regulates synaptic vesicle docking and priming. Nat Commun 8, 15293 (2017).

79. Gracheva, E. O., Hadwiger, G., Nonet, M. L. & Richmond, J. E. Direct interactions between C. elegans RAB-3 and Rim provide a mechanism to target vesicles to the presynaptic density. Neurosci Lett 444, 137–142 (2008).

80. Kaeser, P. S. et al. ELKS2alpha/CAST deletion selectively increases neurotransmitter release at inhibitory synapses. Neuron 64, 227–39 (2009).

81. Lu, J. et al. Solution structure of the RIM1α PDZ domain in complex with an ELKS1b C- terminal peptide. J Mol Biol 352, 455–466 (2005).

82. Chen, L. Y., Jiang, M., Zhang, B., Gokce, O. & Sudhof, T. C. Conditional Deletion of All Neurexins Defines Diversity of Essential Synaptic Organizer Functions for Neurexins. Neuron 94, 611–625.e4 (2017).

83. Südhof, T. C. Synaptic Neurexin Complexes: A Molecular Code for the Logic of Neural Circuits. Cell 171, 745–769 (2017).

84. Luo, F., Sclip, A., Jiang, M. & Südhof, T. C. Neurexins cluster Ca2+ channels within the presynaptic active zone. EMBO J 39, e103208 (2020).

85. Missler, M. et al. Alpha-neurexins couple Ca2+ channels to synaptic vesicle exocytosis. Nature 424, 939–48. (2003).

86. Oh, K. H., Krout, M. D., Richmond, J. E. & Kim, H. UNC-2 CaV2 Channel Localization at Presynaptic Active Zones Depends on UNC-10/RIM and SYD-2/Liprin-α in Caenorhabditis elegans. J Neurosci 41, 4782–4794 (2021).

87. Kaeser, P. S. et al. RIM1alpha and RIM1beta are synthesized from distinct promoters of the RIM1 gene to mediate differential but overlapping synaptic functions. J Neurosci 28, 13435–13447 (2008).

88. Dymecki, S. M. Flp recombinase promotes site-specific DNA recombination in embryonic stem cells and transgenic mice. Proc Natl Acad Sci U S A 93, 6191–6196 (1996).

89. Han, K. A. et al. PTPσ Drives Excitatory Presynaptic Assembly via Various Extracellular and Intracellular Mechanisms. The Journal of Neuroscience 38, 6700–6721 (2018).

90. Strauss, S. & Jungmann, R. Up to 100-fold speed-up and multiplexing in optimized DNA- PAINT. Nat Methods 17, 789–791 (2020).

91. Chen, J.-H., Blanpied, T. A. & Tang, A.-H. Quantification of trans-synaptic protein alignment: A data analysis case for single-molecule localization microscopy. Methods 174, 72–80 (2020).

92. Zürner, M., Mittelstaedt, T., tom Dieck, S., Becker, A. & Schoch, S. Analyses of the spatiotemporal expression and subcellular localization of liprin-α proteins. J Comp Neurol 519, 3019–3039 (2011).

93. Zhang, M. et al. Rational design of true monomeric and bright photoactivatable fluorescent proteins. Nat Methods 9, 727–729 (2012).

